# Cytokine-induced nuclear translocation of STAT1 via a non-transferable NLS

**DOI:** 10.64898/2026.09.28.754937

**Authors:** Pramod Kumar, James R. Cavey, Ruoyu Yang, Oran White, Andreas Begitt, Liu He, Stephanie S. Suinn, Nathan F. Bellis, Ravi K. Lokareddy, Uwe Vinkemeier, Gino Cingolani

## Abstract

The targeting function of nuclear localization signals (NLSs) is generally considered independent of a protein’s native sequence or fold and is readily transferable to heterologous cargos. Contrary to this paradigm, rapid nuclear translocation of phosphorylated STAT1 (pSTAT1) following cytokine stimulation requires importin β, Ran–GTP, and the importin α5 isoform, which recognizes a non-transferable NLS. Here, we present cryo-EM structures of pSTAT1 bound to importin α5, revealing an asymmetric 2:1 complex that diverges from canonical NLS-mediated cargo recognition. Importin α5 occupies the DNA-binding groove of the pSTAT1 dimer, with a single STAT1 N-terminal domain positioning the C-terminal Armadillo repeats 9–10 (S1B domain) orthogonal to the DNA-binding interface. This interface is also targeted by the Ebola virus protein VP24, an antagonist of interferon signaling. We further show that Ran–GTP alone is insufficient to trigger nuclear release of pSTAT1, which additionally requires the exportin CAS. A cryo-EM reconstruction of the CAS–Ran–GTP–α5 complex, supported by in vitro competition assays, demonstrates that CAS and pSTAT1 are mutually exclusive ligands for importin α5. Together, these findings define the molecular choreography of cytokine-induced STAT1 nuclear translocation and release, establishing a general paradigm for STAT family signaling.

## INTRODUCTION

Cytokine signaling through the Janus Kinase (JAK)–Signal Transducer and Activator of Transcription (STAT) pathway converts extracellular immune stimuli into transcriptional responses that maintain cellular homeostasis and host defense. Among STAT family members, STAT1 is the principal effector of interferon (IFN) signaling, controlling antiviral, antiproliferative, and immunomodulatory gene expression programs ^1^. STAT1 exerts its function through a conserved modular architecture that couples receptor activation to gene transcription ^2^. Biochemical and structural studies have shown that STAT1 consists of three structural units ^3–8^ (**Fig. 1a**). The N-terminal domain (ND, residues ∼1–124) mediates dimerization, tetramerization, and nuclear import. A short linker (residues 125–136) connects ND to the STAT1 core, which contains four tandem domains: a coiled-coil domain (CCD, residues ∼137–317) that supports dimerization and cofactor interactions; a DNA-binding domain (DBD, residues ∼320–487) recognizing palindromic motifs; a linker domain (LD, residues ∼488–575) stabilizing DNA-bound dimers; and an SH2 domain (residues ∼577–683) that binds phosphotyrosines and includes the activating Tyr701. Beyond the SH2 domain, the unstructured transactivation domain (TAD, residues 713–750) is essential for transcriptional activity and absent in STAT1β.

**Figure 1.**
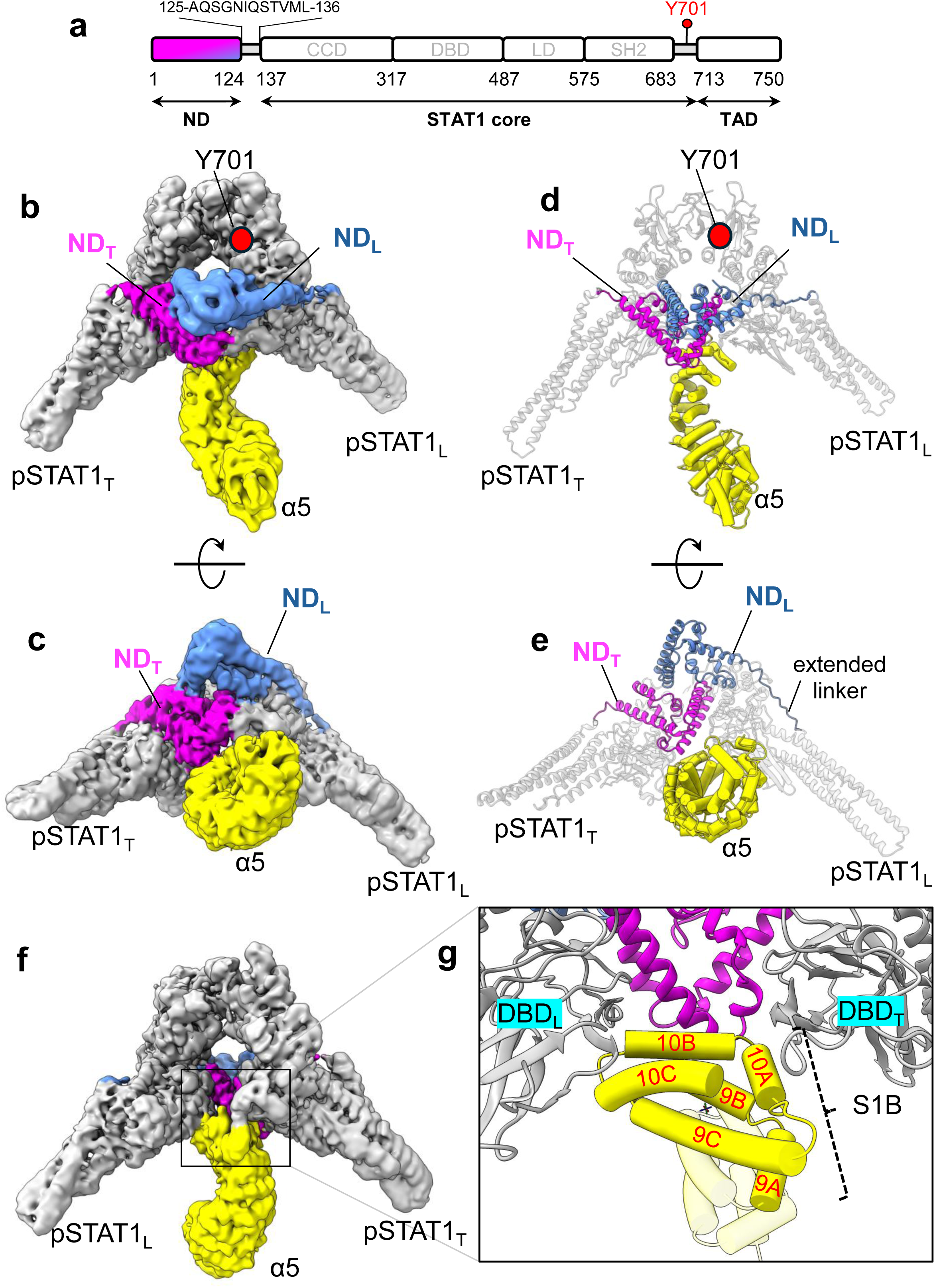
Cryo-EM structure of the 2:1 pSTAT1–α5 complex. (**a**) Schematic diagram of STAT1 domain organization. The amino acid sequence of the flexible linker connecting the ND to the CCD is shown, along with the position of Tyr701. (**b, c**) Cryo-EM density map of the pSTAT1–α5 complex at 3.6 Å resolution, viewed from the N-terminal domain (ND)-facing side (**b**) and after a 90° rotation (**c**). The pSTAT1 core is shown in grey, ND_T_ in magenta, ND_L_ in light blue, and importin α5 in yellow. (**d, e**) Ribbon representations of the fitted pSTAT1–α5 model corresponding to the orientations in (**b**) and (**c**), respectively. (**f, g**) Overall architecture of the pSTAT1–α5 complex with importin α5 facing the viewer; inset in (**g**) shows a close-up of the three-way interaction between importin α5 Arm repeats 9–10 and pSTAT1.

Activation of STAT1 by interferon gamma (IFN-γ) begins at the plasma membrane, where IFN-γ receptor chains assemble with their associated Janus kinases JAK1 and JAK2 ^1^. Ligand engagement triggers receptor trans-phosphorylation, STAT recruitment, and phosphorylation of STAT1 at Tyr701. This single modification drives the formation of parallel pSTAT1 homodimers via reciprocal SH2-phosphotyrosine interactions ^3^, generating a transcriptionally active complex that rapidly accumulates in the nucleus ^9^. Unlike cargos bearing a classical NLS (cNLS) ^10^, pSTAT1 lacks a transferable NLS ^11,12^. Instead, it contains a non-transferable, dimer-specific nuclear-targeting signal in its DNA-binding domain ^5,12^, which is specifically recognized by importin α5 ^11,13,14^. This isoform of the import adaptor importin α has an N-terminal IBB domain ^15^ that recruits importin β ^16,17^ and a core Arm domain that binds cNLS cargo ^18^. The interaction of pSTAT1 with importin α5 is highly selective among importin α isoforms ^19^ and engages both the ND and the DNA-binding domains of pSTAT1 ^20–22^. Nuclear import requires Ran–GTP ^23^, which facilitates translocation of the pSTAT1–α5–β import complex through the nuclear pore complex (NPC). By binding to importin β with high affinity, Ran–GTP triggers an allosteric change that reduces importin β’s avidity for phenylalanine-glycine-rich nucleoporins lining the NPC ^24,25^. Whereas pSTAT1 nuclear accumulation is sustained by continuous nucleocytoplasmic cycling and kinase activity, STAT1 export from the nucleus depends on dephosphorylation by the phosphatase TC45 on Tyr701 ^26^, which is controlled by DNA binding ^27^. Export of unphosphorylated STAT1 by diffusion or active transport completes the signaling cycle ^28^.

Despite a well-defined biochemical framework, the structural principles governing how pSTAT1 dimers are recognized and translocated by importin α5 remain incompletely understood, limiting our ability to mechanistically link plasma membrane activation to nuclear transcriptional activity. Here, we present the atomic structure of full-length pSTAT1 bound to importin α5, revealing the molecular basis for isoform specificity. We identify a previously unrecognized cargo-binding surface within Arm repeats 9–10, which we term the STAT1-binding (S1B) domain. We further show that the importin α5 Arm core (Arm repeats 2–8) and S1B (Arm repeats 9–10) are distinct, non-overlapping interfaces, which can simultaneously engage a cNLS cargo and pSTAT1. Finally, we show that Ran–GTP and the export receptor CAS, also a member of the importin β family ^29^, dissociate the pSTAT1 import complex and identify the key requirements for STAT1-specific nuclear import inhibition.

## RESULTS

### Cryo-EM structures of pSTAT1 bound to importin α5

Single-particle analysis of pSTAT1–α5 complexes identified two main oligomeric states: a homodimer bound to one importin α5 (2:1 assembly), predominant at low micromolar concentrations, and a tetramer bound to two importin α5 molecules (4:2 assembly), abundant at supraphysiological concentrations. We reconstructed both assemblies at 3.6 Å resolution (**Supplementary Figs. 1**–**2** and **Table 1**). The 2:1 complex is consistent with the oligomerization stoichiometry observed in solution at low micromolar concentrations of pSTAT1 and importins, and likely represents the physiological state of the pSTAT1 import complex ^20^. In contrast, the 4:2 assembly, which forms at pSTAT1 concentrations >10 μM, is unlikely to be a physiologically relevant species in the cytoplasm of human cells, where total importin α and STAT1 concentrations have been estimated at ∼1 μM ^30^ (with the α5 isoform likely substantially lower) and ∼40 nM ^31^, respectively.

**Table 1.**
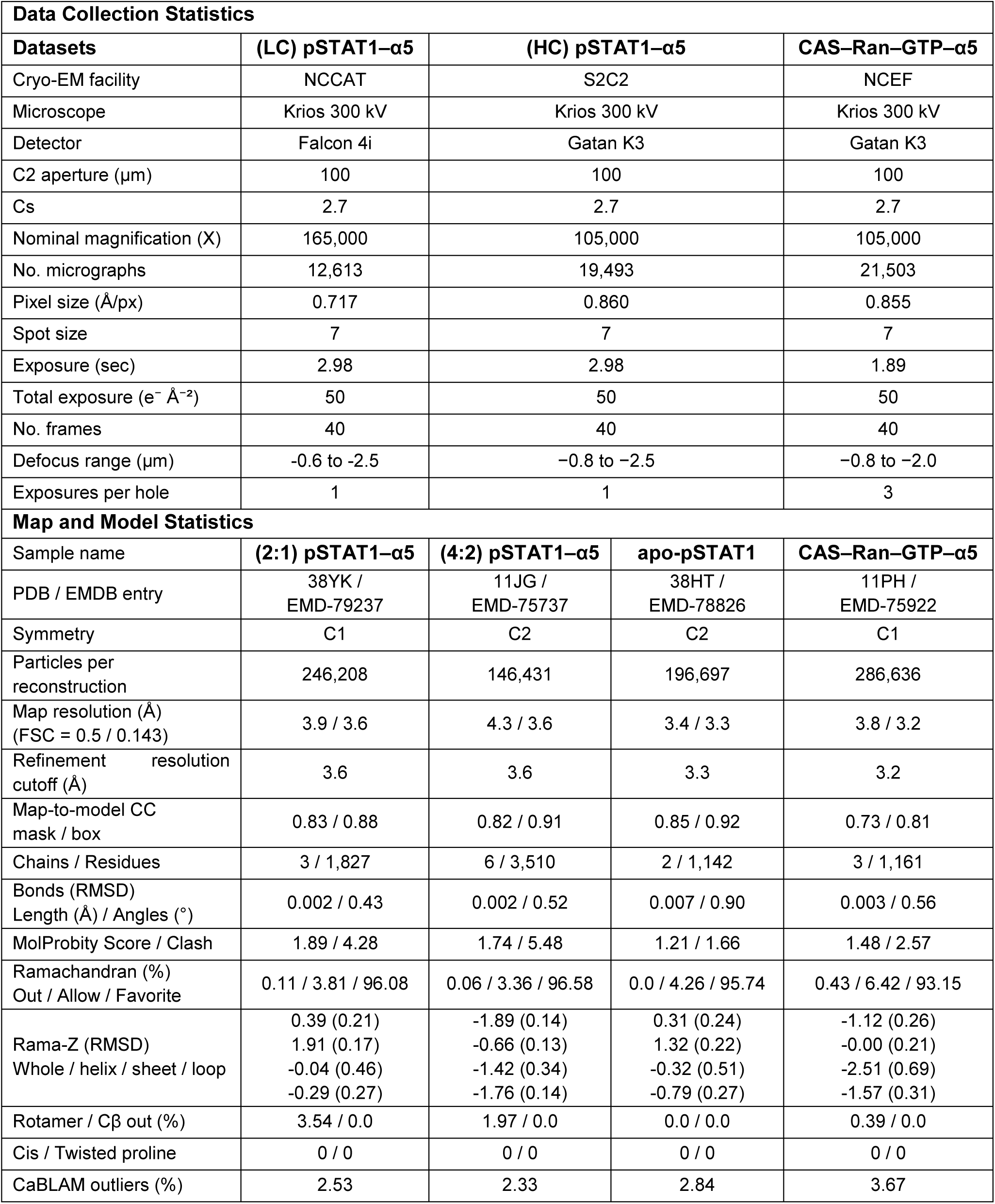
Cryo-EM data collection and model refinement statistics.

The reconstruction of the pSTAT1–α5 2:1 complex, vitrified at ∼1 mg/ml (equivalent to ∼7.5 μM pSTAT1) (**Fig. 1b, c**), revealed a highly asymmetric heterodimer in which the two pSTAT1 protomers adopt distinct conformational states defined by their NDs. The cryo-EM density enabled us to build a complete atomic model of dimeric pSTAT1 (residues 2–717) bound to importin α5 (residues 86–507), lacking the IBB domain (**Fig. 1d, e**). All 10 Arm repeats of importin α5 are visible in the reconstruction, highlighting its rigidity ^18^. Asymmetry is conferred by the NDs, which form a handshake interface ^32–34^, and by the linkers (residues 125–136) that connect them to the pSTAT1 core (**Fig. 1a**). In one protomer (pSTAT1_T_), the ND adopts a “tight” conformation (ND_T_), stabilized by a well-resolved linker and a direct interaction with importin α5 (**Fig. 1e**). In the second protomer (pSTAT1_L_), the ND adopts a “loose” conformation (ND_L_), in which it is displaced from the core and connected to the beginning of the CCD helix by an extended linker (**Fig. 1e**). Despite these differences in ND positioning, both pSTAT1 cores maintain similar tertiary structures (root-mean-square deviation, RMSD = 0.53 Å), with variability confined to the ND-linker region. Importin α5 binds pSTAT1 exclusively through Arm repeats 9– 10, which we designate as the STAT1-binding (S1B) domain (**Fig. 1f**–**g**). The S1B domain engages the DNA-binding domains of both protomers, as well as the ND of the tight protomer (ND_T_), forming a tripartite binding interface (**Fig. 1g**).

### ND swapping stabilizes a tetrameric assembly of pSTAT1 and importin α5

A second dataset collected from grids vitrified at a higher concentration of the pSTAT1–α5 complex (∼1.5 mg/ml, equivalent to 11.5 μM pSTAT1) yielded a 3.6 Å resolution reconstruction of a much larger 4:2 pSTAT1–α5 assembly (**Fig. 2a**–**b**). This larger assembly contains two pSTAT1–α5 complexes arranged as a dimer of heterodimers (**Fig. 2c**–**d**). Each pSTAT1 dimer adopts a similar overall organization in the 4:2 and 2:1 complexes (RMSD ∼1.5 Å), but the tetrameric assembly is stabilized by ND-swapping. We designated each homodimer as primary (protomers a, b) and secondary (protomers a′, b′) (**Fig. 2e**). They interlock through ND swapping, yielding a tetrameric topology stabilized by pairs of ND_T_–ND_L_′ and ND_L_–ND_T_′ “handshake” interfaces. In contrast, the orientation of importin α5 differs slightly between the two assemblies, with α5 adopting a spoon-like orientation in the 4:2 complex compared with the more orthogonal orientation observed in the 2:1 complex (**Fig. 2f**).

**Figure 2.**
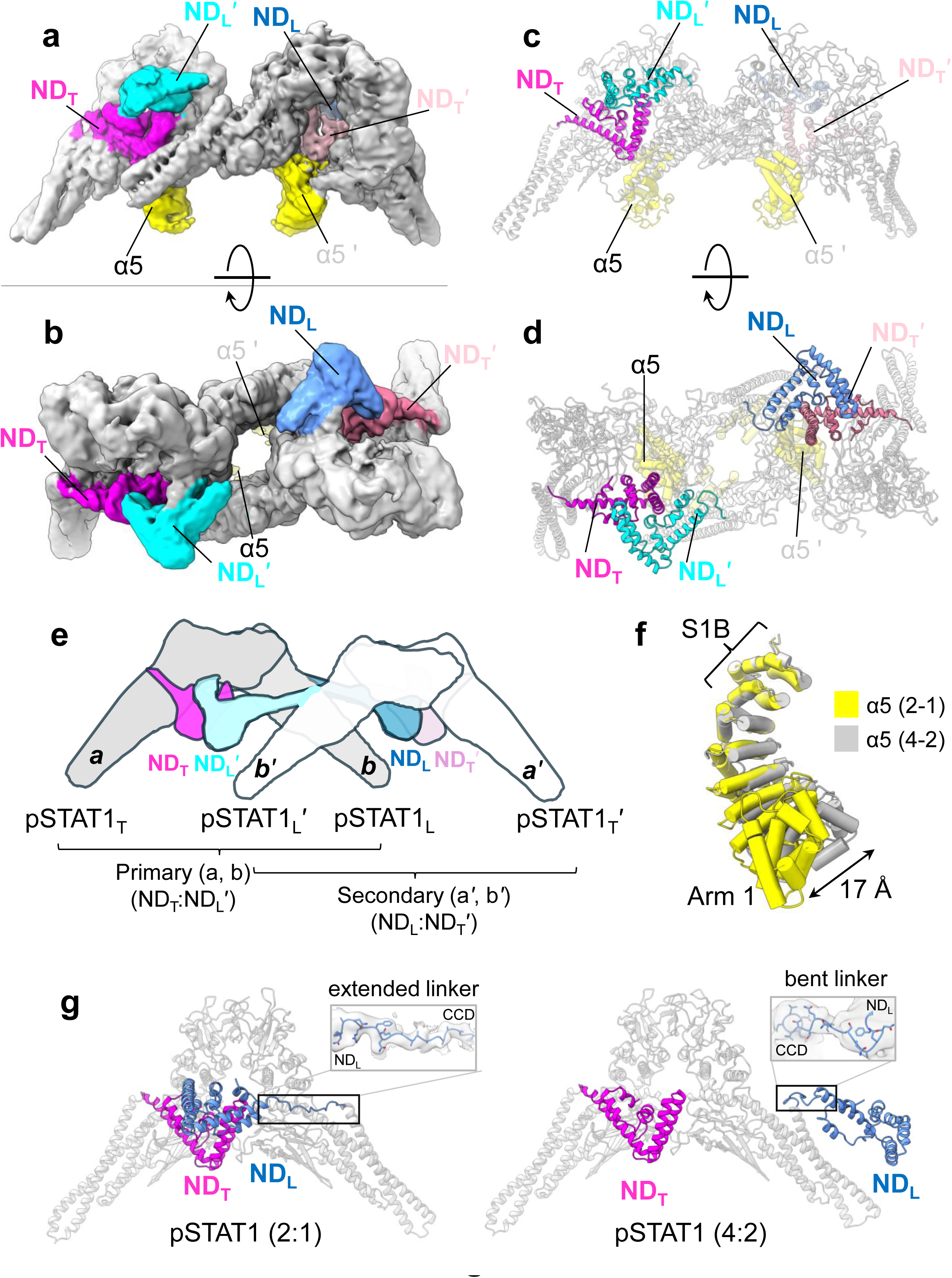
Cryo-EM structure of the ND-swapped 4:2 pSTAT1–α5 complex. (**a, b**) Cryo-EM density map of the tetrameric pSTAT1–α5 complex shown from the ND-facing side (**a**) and a 90° rotated view (**b**). The C2-symmetrized cryo-EM density reached a global resolution of 3.6 Å, based on the FSC = 0.143 criterion. (**c, d**) Ribbon representations of the ND-swapped pSTAT1– α5 models corresponding to the orientations in panels (**a, b**). (**e**) Cartoon representation of the pSTAT1 tetramer organization (importin α5 is not shown). pSTAT1 dimers, formed by protomers (a, b) and (a′, b′), are colored gray and white, respectively; NDs are colored magenta (ND_T_) and cyan (ND_L_′) for protomers (a, b′), and pink (ND_T_′) and blue (ND_L_) for protomers (a′, b). (**f**) Superimposition of importin α5 from the 2:1 (yellow) and 4:2 (gray) assemblies reveals a distinct orientation of ARM repeats 1–6. The maximum displacement of Arm 1 in the two states is about 17 Å. (**g**) Comparison of the closed pSTAT1 dimer in the 2:1 assembly (left) and the open pSTAT1 dimer in the 4:2 assembly (right). The linker connecting ND_L_ to the coiled-coil domain of protomer *b* is extended in the 2:1 assembly and bent in the 4:2 assembly. Zoom-in panels show the cryo-EM density of the linkers overlaid on the refined model.

Comparison of the pSTAT1 dimers formed by protomers a and b in the 2:1 and 4:2 assemblies reveals that ND swapping is enabled by a conformational change in the linker connecting ND_L_ to the coiled-coil domain (**Fig. 2g**). This linker is fully extended in the 2:1 assembly, spanning ∼44 Å, whereas it adopts a partially bent conformation in the domain-swapped 4:2 assembly, spanning about 30 Å. Importin α5 binding remains asymmetric: only ND_T_ engages α5, while ND_L_ is exchanged between the two pSTAT1 dimers. Tetramerization of pSTAT1 dimers is a concentration-dependent process that occurs spontaneously in solution above 1 μM ^20,31^ and potentially at lower concentrations on DNA, through cooperative binding of adjacent pSTAT1 dimers to tandem GAS sites ^35^.

In addition to the well-defined 4:2 pSTAT1–α5 assembly (**Fig. 2a–d**), the dataset from grids vitrified at a higher concentration also contained heterogeneous intermediate conformations (**Supplementary Fig. 3a**–**c**). These included pSTAT1–α5 heterodimers (a, b) associated with a partially occupied second heterodimer (a′, b′), often with missing density for protomer b′. These intermediates likely represent domain-swapped tetramers in which a single ND bridges two pSTAT1 dimers, generating unstable assemblies and diffuse density for the second dimer.

### Structural basis for importin α5 specificity

The cryo-EM structure of the pSTAT1–α5 complex delineates the molecular basis for isoform-specific recognition. Binding to the S1B domain involves contributions from both protomers, forming an extensive, highly asymmetric interface (**Fig. 1f**). The tight protomer provides a major contact surface (587 Å²) through its ND_T_ and DBD_T_, forming a wrench-head–like interaction. The loose protomer contributes a smaller interface (448 Å²), restricted to engagement of its DBD_L_. Overall, pSTAT1 engages residues from all three helices of Arm 10 and helix B of Arm 9 through four salt bridges, four hydrogen bonds, and 23 van der Waals contacts (≤ 4.0 Å) (**Fig. 3a** and **Supplementary Table 1**). Consistent with these extensive interactions, pSTAT1 binds importin α5 *in vitro* with a dissociation constant of K_d_ = 191 ± 20 nM ^20^. The stability of the binding interface is primarily maintained by hydrophobic packing, supported by specific polar interactions. These include two salt bridges involving the pSTAT1_T_ DBD (DBD_T_) residues; two salt bridges and two hydrogen bonds from residues in the ND_T_ and two hydrogen bonds involving the pSTAT1_L_ DBD (DBD_L_) (**Fig. 3a**). As previously noted, STAT1 ND_L_ does not make direct contacts with importin α5 and remains highly dynamic.

**Figure 3.**
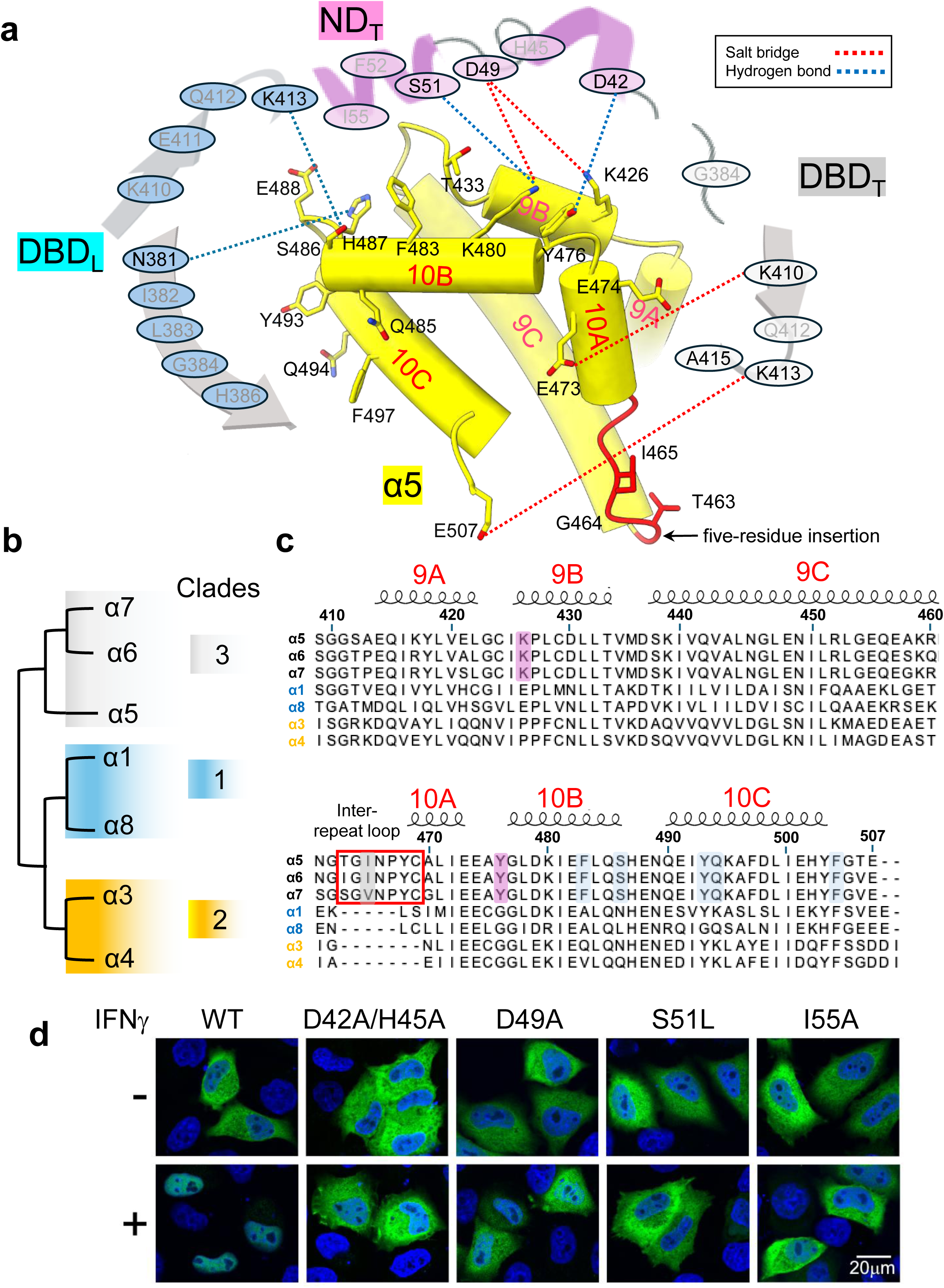
Determinants of importin α5 specificity for pSTAT1. (**a**) Schematic of the importin α5 S1B domain (yellow helices) showing the tripartite pSTAT1 recognition interface, with DBD_L_ (light blue), ND_T_ (magenta), and DBD_T_ (light gray). All pSTAT1 residues within ≤ 4.0 Å of the S1B domain are shown schematically. Dashed lines denote salt bridges (red) and hydrogen bonds (blue). (**b**) Phylogenetic relationships among human importin α isoforms are displayed as a cladogram. Importin α5 clusters within clade 3 together with α6 and α7. (**c**) Sequence alignment of Arm repeats 9–10 from all importin α isoforms. Clade 3 members are at the top. Red box highlights the insertion loop characteristic of clade 3. α5 residues contacting ND_T_ are magenta; those contacting DBD_L_ are light blue. (**d**) Confocal fluorescence micrographs of HeLa cells expressing GFP-tagged WT STAT1 or point mutants before and after treatment with 50 U mL^-1^ IFN-γ for 1 h. Nuclear chromatin is stained with Hoechst dye (blue).

Structural and bioinformatic analyses reveal three key features in importin α5 Arm repeats 9–10 that underlie isoform-specific recognition of pSTAT1. <u>First</u>, among the three major clades (1, 2 and 3), importin α5 belongs to clade 3, along with isoforms α6 and α7 ^19^ (**Fig. 3b**). In these isoforms, insertions of five and seven residues, relative to clade 1 and clade 2 isoforms, respectively, extend the inter-repeat loop between Arm repeats 9–10 (highlighted in red in **Fig. 3a** and outlined in red in **Fig. 3c**), creating a surface that fits snugly against the pSTAT1 DNA-binding groove. Ile465 of importin α5, exposed on this insertion loop, contacts Gln412 and Lys413 in the DBD_T_ (**Supplementary Table 1**), residues critical for nuclear import and DNA-binding ^5,11^. <u>Second</u>, a limited number of isoform-specific residues in importin α5 make unique contacts with pSTAT1. In Arm 9 (**Fig. 3c**), Lys426 forms a salt bridge with pSTAT1 ND_T_ Asp49, whereas Tyr476, located in the loop connecting Arm 10 helices A and B, engages ND_T_ His45, Asp42, and Phe52 (**Supplementary Table 1**). In Arm 10 (**Fig. 3c**, highlighted in light blue), Ser486 forms a hydrogen bond with Lys413 in DBD_L_, while Tyr493, Gln494, and Phe497 on helix C establish additional van der Waals contacts with Asn381, Ile382, Leu383, His386, and Gly384 of DBD_L_ (**Supplementary Table 1**). <u>Finally</u>, the interface between the pSTAT1 ND_T_ and importin α5 is stabilized by a network of electrostatic and hydrogen-bonding interactions. These include two salt bridges between ND_T_ Asp49 and importin α5 Lys426 and Lys480 (Arm repeats 9–10); hydrogen bonds between ND_T_ Asp42 and Tyr476, as well as ND_T_ Ser51 and Lys480 across helices 9B and 10B (**Fig. 3a** and **Supplementary Table 1**). Notably, Lys480 of importin α5 uniquely bridges three ND_T_ residues, Asp49, Ser51, and Ile55, highlighting its central role in conferring binding specificity.

To validate our structural analysis, we performed a cell-based assay using pSTAT1 variants carrying substitutions at key ND_T_ residues Ile55, Ser51, Asp49, His45, and Asp42 (**Fig. 3d**). Mutations disrupting the N-domain interface (Asp42Ala/His45Ala, Asp49Ala, Ser51Leu, and Ile55Ala) markedly impaired IFN-γ-inducible nuclear translocation (**Fig. 3d**), despite normal Tyr701 phosphorylation upon IFN-γ stimulation (**Supplementary Fig. 4**). These results confirm the critical role of the ND_T_–S1B interaction in mediating pSTAT1 nuclear import in living cells.

### Overlapping yet distinct modes of pSTAT1 interaction with importin α5 and DNA

The higher-concentration dataset also yielded a 3.3 Å resolution reconstruction of apo-pSTAT1 (**Fig. 4a** and **Supplementary Fig. 2**), which, despite lacking density for the NDs, revealed a rather open conformation of the DBDs, similar to that observed with importin α5 (RMSD 1.5 Å). We next compared the three states of pSTAT1, namely apo, bound to importin α5, and DNA. Strikingly, the importin α5 binding surface on pSTAT1 partially overlaps its DNA-binding interface (PDB ID: 1BF5) ^3^ (**Fig. 4b, c**). Superposition of individual pSTAT1 cores lacking the N-terminal domain from the 2:1 complex onto their DNA-bound counterparts yields an RMSD of 1.2 Å, indicating strong conservation of the monomeric core fold. Despite this similarity, the quaternary architecture of the pSTAT1 dimer differs substantially between the importin α5-bound and DNA-bound states, corresponding to cytoplasmic and nuclear binding partners, respectively. Both importin α5 and double-stranded DNA (dsDNA) engage the dimeric interface of pSTAT1 (**Fig. 4b, c**), but their relative orientations differ markedly. DNA runs perpendicular to the pSTAT1 twofold axis, whereas the superhelical axis of importin α5 aligns parallel to it, rendering the two substrates essentially orthogonal. DNA occupies the full depth of the pSTAT1 groove, positioned ∼30 Å above importin α5, whereas importin α5 interacts only with the entrance of the DNA-binding groove, forming an asymmetric, symmetry-mismatched interface.

**Figure 4.**
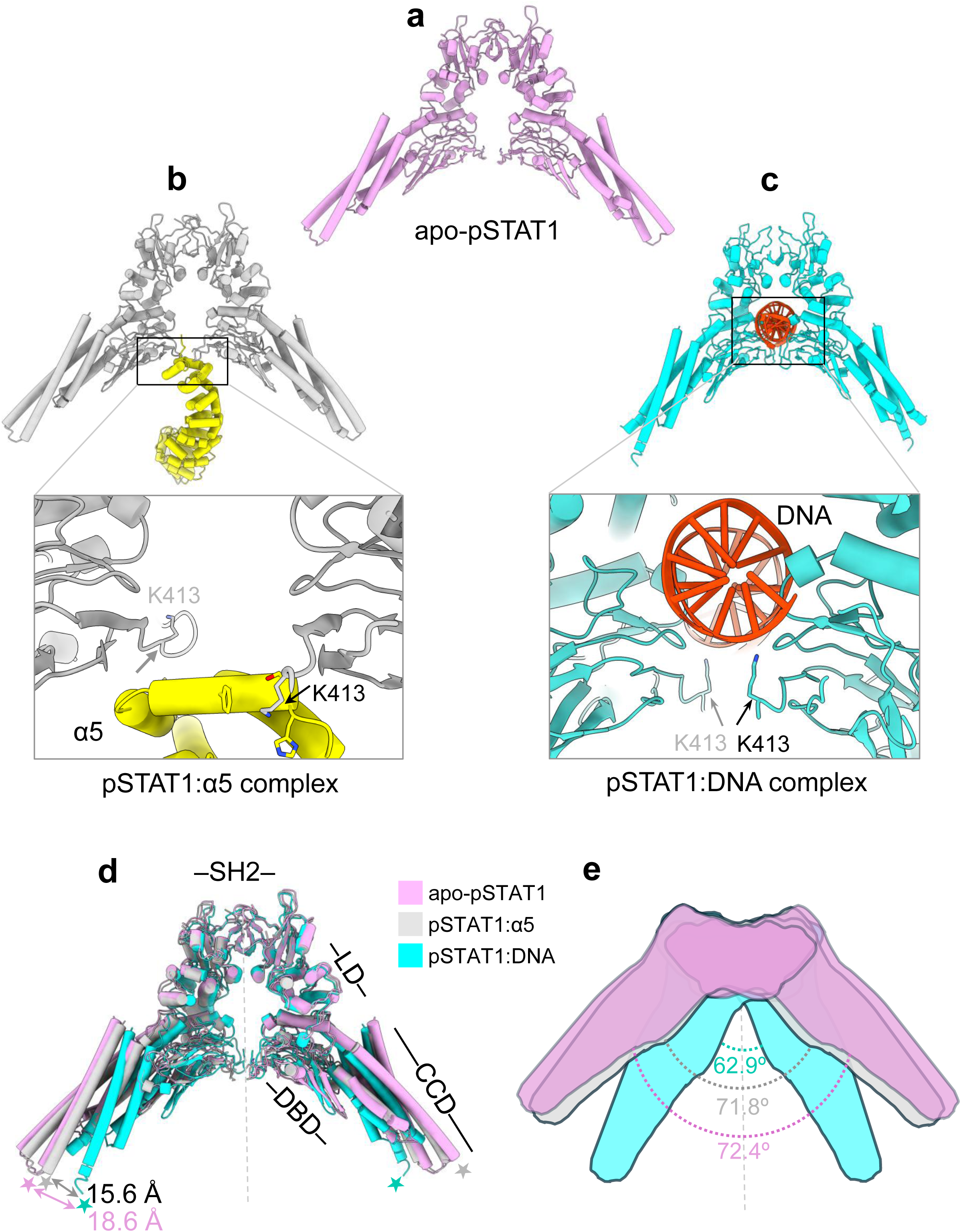
Distinct binding modes of importin α5 and dsDNA for pSTAT1. (**a**) Ribbon diagram of apo-pSTAT1 (light magenta), refined to 3.3 Å resolution. The NDs are not resolved in the reconstruction. (**b**) Ribbon diagram of the pSTAT1 core (lacking NDs) (gray) bound to importin α5 (yellow) presented in this paper. The NDs are not shown in the ribbon diagram. (**c**) Ribbon diagram of pSTAT1 core (cyan) bound to DNA (orange) from PDB: 1BF5. The inset panels in (**b, c**) show a detailed view of importin α5 Arm 10 and DNA, as well as the role of pSTAT1 Lys413 in their interaction. (**d**) Superimposition of pSTAT1 cores in apo-pSTAT1 (light magenta) and bound to importin α5 (gray) and DNA (cyan), showing maximum displacements of 18.6 Å at the coiled-coil tip between apo- and DNA-bound pSTAT1 and of 15.6 Å between α5- and DNA- bound pSTAT1. (**e**) Schematic comparison of the angles between pSTAT1 protomers in apo-pSTAT1 (72.4°) and in the importin α5-bound (71.8°) and DNA-bound (62.9°) states.

Although they engage distinct molecular surfaces, importin α5 and DNA share a critical contact site: Lys413 of STAT1. In all pSTAT1–DNA complexes (PDB IDs: 1BF5, 8YYV, and 8YYU ^3,35^), Lys413 projects inward from the β-barrel to make electrostatic contacts with the DNA phosphate backbone (**Fig. 4c**). In contrast, in the importin α5-bound tight protomer, Lys413 adopts an outward-facing orientation (**Fig. 4b**). In the tight protomer, it anchors into the interface with importin α5 residues Glu507 and Ile465, whereas in the loose protomer, it engages Ser486, Phe483, and His487 (**Fig. 3a** and **Supplementary Table 1**). This conformational switch highlights Lys413’s dual role as a hinge residue that toggles between DNA- and importin-binding modes. Consistent with this model, mutation of residue Lys413 abolishes both DNA binding and nuclear import of pSTAT1 ^5,11^.

Importin α5 and DNA each induce global conformational rearrangements in pSTAT1. Compared with apo-pSTAT1, which adopts the most open conformation, binding to either DNA or importin α5 induces a more compact quaternary structure. Superposition of the DNA- and importin α5-bound dimeric assemblies yields a global RMSD of 4.7 Å. Relative to apo-pSTAT1, the largest displacement occurs at the coiled-coil tip, reaching 18.6 Å in the DNA-bound state; the corresponding displacement between the importin α5- and DNA-bound states is 15.6 Å (**Fig. 4d**). This transition is accompanied by a closure of the inter-protomer angle from 72.4° in the apo-pSTAT1 to 71.8° in the importin α5-bound state, with a more pronounced closure to 62.9° upon DNA binding (**Fig. 4e**). Together, these changes reflect conformational adaptation of the CCD and DBDs that accommodates importin α5 binding on a more splayed surface, orthogonal to the DNA-binding plane.

### pSTAT1 and NLS cargo can bind simultaneously to importin α5

Structural analysis identifies importin α5 Arm repeats 9–10 as a spatially segregated platform for pSTAT1 engagement. The S1B domain (**Fig. 5a**) is distinct from the major groove (Arm repeats 2–4) and the minor groove (Arm repeats 6–8), which are bound by the N-terminal IBB domain ^15,36^ (**Fig. 5b**) and cNLS-containing cargo ^10,37^ (**Fig. 5c**). In the reconstruction (**Fig. 1b, d**), both the major and minor NLS-binding sites are unoccupied and potentially available to bind NLS cargos. To test whether importin α5 can simultaneously accommodate pSTAT1 and an NLS cargo (**Fig. 5d**), we assembled dimeric and trimeric complexes of importin α5 *in vitro* using purified proteins (**Supplementary Fig. 5a**). As NLS cargo, we used a protein derived from Influenza Virus Polymerase subunit PB2, previously shown to be specific to importin α5 ^18^, fused to an N-terminal histidine tag (H-NLS). Native gel electrophoresis revealed progressive migration shifts consistent with the formation of a ternary pSTAT1–α5–H-NLS complex (**Fig. 5e**, lanes 6– 8), which migrated above the heterodimeric pSTAT1–α5 (**Fig. 5e**, lane 5) but below free pSTAT1 (**Fig. 5e**, lanes 2, 9). Immunoblotting of PVDF transfers with anti-His antibodies confirmed the presence of H-NLS in both the heterodimeric importin α5–H-NLS (**Fig. 5f**, lane 4) and ternary pSTAT1–α5–H-NLS complexes (**Fig. 5f**, lanes 6–8). These data indicate that *in vitro* importin α5 can simultaneously bind pSTAT1 and a cNLS cargo, suggesting that nuclear import of pSTAT1 can occur alongside that of cargos containing cNLS.

**Figure 5.**
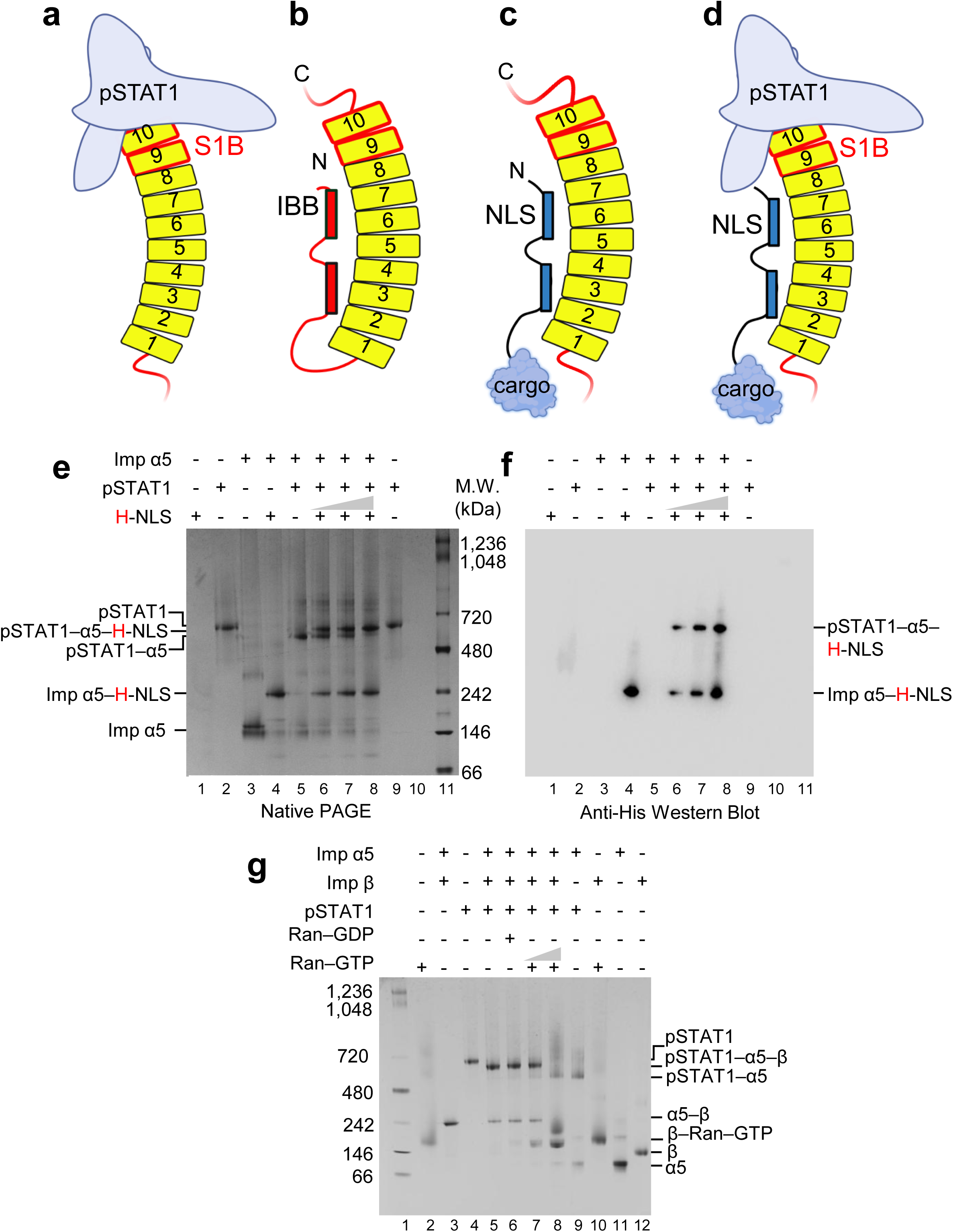
pSTAT1 and cNLS cargo bind distinct sites on importin α5. (**a–d**). Schematics of importin α5 bound to pSTAT1 (**a**), its N-terminal IBB domain (**b**), a bipartite cNLS (**c**), or both pSTAT1 and cNLS (**d**). (**e**) Native gel electrophoresis showing that sequential addition of His-PB2-NLS (H-NLS) to a preformed pSTAT1–α5 complex (lanes 6–8) yields a distinct species migrating between free pSTAT1 (shown in lanes 2 and 9) and the pSTAT1–α5 complex (lane 5), consistent with formation of a ternary pSTAT1–α5–H-NLS assembly. (**f**) Immunoblot of the native gel in (**e**) using an anti-His antibody confirms incorporation of H-NLS into both the ternary pSTAT1–α5–H-NLS complex and the importin α5–H-NLS heterodimer (lanes 6–8). (**g**) Native gel analysis shows that Ran–GTP (lanes 7–8), but not Ran–GDP (lane 6), displaces importin β (lane 12) from the pSTAT1–α5–β complex (lane 5). This reaction yields two heterodimeric species (lane 8): pSTAT1–α5 and importin β–Ran–GTP, which co-migrate with the respective controls in lanes 9 and 10.

### Ran–GTP releases importin β but not pSTAT1 from importin α5

Because the pSTAT1- and cNLS-binding sites on importin α5 do not overlap, we next examined how the pSTAT1 import complex dissociates upon nuclear entry. For cNLS cargo, Ran–GTP is necessary and sufficient to disassemble the import complex, displacing importin α from importin β ^38,39^ and releasing the cargo through IBB-mediated autoinhibition ^36^ (**Fig. 5b**). This mechanism, originally characterized in importin α1, is conserved in importin α5, which exhibits weaker autoinhibition ^18^. Using purified proteins (**Supplementary Fig. 5b**), we first formed a stable pSTAT1–α5–β complex (**Fig. 5g**, lane 5), which was not disrupted upon addition of Ran–GDP (**Fig. 5g**, lane 6). In contrast, titrating in Ran–GTP dissociated importin β from the pSTAT1–α5– β complex, while the pSTAT1–α5 complex remained intact (**Fig. 5g**, lanes 7–8). The persistence of the pSTAT1–α5 heterodimer likely reflects structural constraints, as the IBB domain cannot reach Arm repeats 9–10 to displace pSTAT1 ^36^. These results indicate that Ran–GTP selectively releases importin β from importin α5 without freeing pSTAT1, suggesting that an additional cellular factor is needed to fully disassemble the pSTAT1 import complex and release the activated transcription factor into the nucleus.

### Ebola VP24 and Nup50 bind the C-terminus of importin α5

To investigate how pSTAT1 is released from importin α5 in the nucleus, we examined known importin α5-binding proteins implicated in the disassembly of import complexes. Previous crystal structures have identified two substrates that bind the C-terminus of importin α5, or the closely related isoform α6, in a manner analogous to pSTAT1: the nucleoporin Nup50 (residues 1–109), which engages both the minor NLS-binding pocket and Arm 10 of importin α5 ^40^ (**Fig. 6a**), and the Ebola virus VP24 ^41^ (**Fig. 6b**), which was crystallized bound to the clade 3 isoform importin α6 (**Fig. 3b**). Structural modeling indicates that Nup50 and VP24 bind overlapping surfaces within the S1B domain, suggesting that this region serves as a shared recognition platform for host and viral cargos. Within this interface, Nup50 contacts importin α at Glu473, His487, and Ser486 (**Fig. 6a**), all of which are implicated in pSTAT1 binding (**Fig. 3a** and **Supplementary Table 1**), and also engages the minor NLS ^40^. Similarly, VP24 contacts importin α6 at Glu474 and Lys481 (**Fig. 6b**), equivalent to importin α5 Glu473 and Lys480, two key determinants of pSTAT1 recognition (**Fig. 3a** and **Supplementary Table 1**). Despite these structural similarities, VP24 and Nup50 differ markedly in function. VP24 is a potent inhibitor of pSTAT1 nuclear import that blocks the antiviral interferon response ^41,42^, whereas Nup50 is a nucleoporin associated with the NPC basket ^43^ that is thought to promote nuclear import by facilitating cargo release from importin α ^44^. To test whether Nup50 can directly displace pSTAT1 from importin α5, we performed native gel electrophoresis with purified proteins (**Fig. 6c** and **Supplementary Fig. 5c**). We first assembled homogeneous complexes of Nup50–α5 (**Fig. 6c**, lane 4) and pSTAT1– α5 (**Fig. 6c**, lane 5), then gradually added Nup50 into the preassembled pSTAT1–α5 complex (**Fig. 6c**, lanes 6–8). This led to the disappearance of the pSTAT1–α5 species and the emergence of Nup50–α5, along with free pSTAT1 migrating as in the control (**Fig. 6c**, lanes 4 and 9). These results indicate that Nup50 can directly displace pSTAT1 from importin α5 *in vitro*, most likely by competing for overlapping residues within the S1B domain.

**Figure 6.**
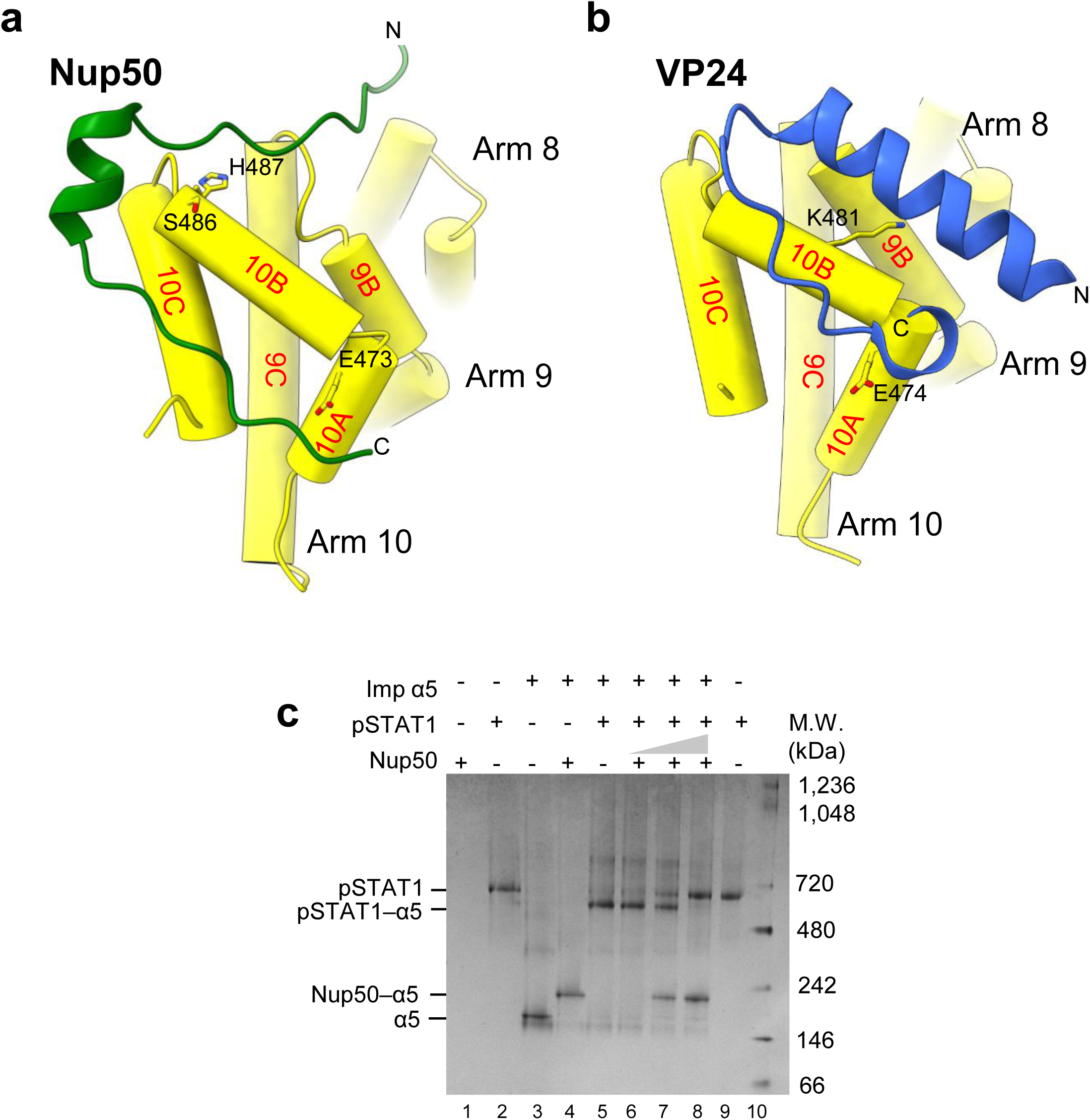
Ebola VP24 and Nup50 engage the importin α5 S1B domain. (**a**) Close-up view of the interaction between the C-terminus of importin α5 (yellow) and Nup50 (residues 20–48, green; PDB 3TJ3). (**b**) Close-up view of the interaction between the C-terminus of importin α6 (yellow) and Ebola virus VP24 (residues 113–141, blue; PDB 4U2X). In (**a, b**), the highlighted side chains correspond to key pSTAT1-binding determinants of importin α5 or α6 engaged by Nup50 or VP24. (**c**) Native gel electrophoresis showing that addition of Nup50 (lane 1) to a preformed pSTAT1–α5 complex (lanes 6–8) displaces pSTAT1, yielding free pSTAT1 and a Nup50–α5 complex. Controls for free pSTAT1 (lanes 2 and 9) and the Nup50–α5 complex (lane 4) are shown.

### Nup50 does not regulate STAT1 nuclear import under physiological conditions

To assess the role of Nup50 in pSTAT1 nuclear translocation in living cells, we generated a human 293T cell line using CRISPR-Cas9, expressing a Nup50 variant bearing an internal 28-amino acid deletion encompassing the importin α5-binding region (residues 24–51) (**Supplementary Fig. 6a, b**). Immunoprecipitation with FLAG-tagged importin α5 confirmed that wild-type (WT) Nup50, but not the deletion mutant, interacted with importin α5 (**Supplementary Fig. 7**, top panel).

IFN-γ–induced nuclear accumulation of endogenous STAT1 was comparable in parental WT and Nup50-mutant cells, with similar kinetics and extent within 30 min of stimulation, irrespective of Nup50 binding to importin α5 (**Fig. 7a, b**). To probe potential downstream effects of impaired cargo release, we monitored pSTAT1 dephosphorylation in the nucleus. We reasoned that defective dissociation from importin α5 would stabilize STAT1 dimers in the parallel conformation, resulting in delayed tyrosine dephosphorylation and prolonged nuclear retention ^27,45^. In both WT and Nup50-mutant cells, STAT1 tyrosine phosphorylation decayed within ∼20 min after addition of the kinase inhibitor staurosporine, and nuclear STAT1 levels collapsed after ∼40 min, with no detectable differences between the two cell lines (**Fig. 7c, d**). These data indicate that disruption of the Nup50:importin α5 interaction does not measurably affect STAT1 nuclear import or release under physiological conditions. Thus, Nup50-mediated disassembly of the importin α5–pSTAT1 complex observed *in vitro* may not occur under physiological conditions.

**Figure 7.**
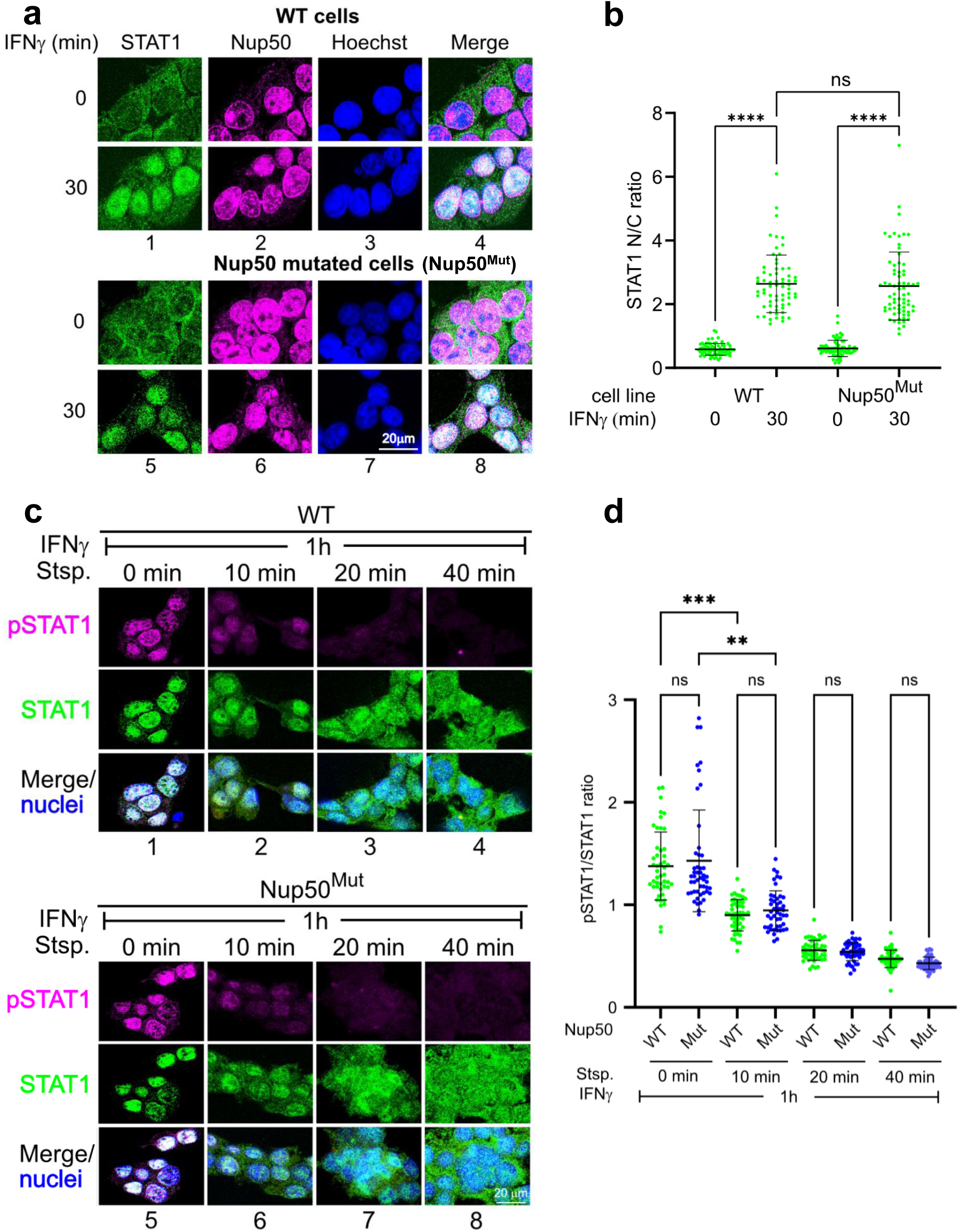
Nuclear import of pSTAT1 in gene-edited cells expressing Δ24–51 Nup50 (Nup50^Mut^). (**a**) Immunocytochemistry results using 293T WT cells (top panels) or gene-edited cells expressing Δ24–51 Nup50 (bottom panels) before and after treatment with IFN-γ. Fixed cells were probed with antibodies against STAT1 (panels 1 and 5) and Nup50 (panels 2 and 6). Nuclei were stained with Hoechst dye (panels 3 and 7). (**b**) Quantification of STAT1 nuclear accumulation (STAT1 nuclear/cytoplasmic fluorescence ratios) of cells shown in (**a**). ∗∗∗∗p < 0.0001 as determined by a Kruskal–Wallis test; ns, not significant. (**c**) Immunocytochemistry micrographs depicting the time course of Tyr701-phosphorylation (top row) and subcellular distribution (middle row) of endogenous STAT1 in WT (rows 1–4) and Δ24–51 Nup50 mutant (rows 5–8) 293T cells. After inducing STAT1 phosphorylation and nuclear accumulation (1 h of IFN-γ), the cells were treated for the indicated times with the kinase inhibitor staurosporine (Stsp) to assess dephosphorylation activity. The merged images additionally show nuclear chromatin (blue; Hoechst stain). (**d**) Quantification of STAT1 dephosphorylation time course (pSTAT1/STAT1 fluorescence ratios) shown in (**c**). ∗∗∗p < 0.001, ∗∗p < 0.01 as determined by a Kruskal–Wallis test; ns, not significant.

### Nuclear import inhibition of pSTAT1/pSTAT2 requires competition for importin α5 and cytoplasmic localization

To assess competition between Ebola VP24 and importin α5, we transfected HeLa cells with mCherry-VP24 and observed strong cytoplasmic retention of pSTAT1, consistent with its established role as an inhibitor of STAT1 nuclear import (**Fig. 8a**) ^41,42^. Notably, VP24 also retained pSTAT2 in the cytoplasm, suggesting that the importin α5-dependent nuclear import mechanism defined here for pSTAT1 extends to STAT2 (**Fig. 8a**). This was corroborated by co-precipitation of activated STAT2, which enters the nucleus as a heterodimer with STAT1 in response to type I and III interferons ^46^, with WT importin α5, but not with its Tyr476Ala mutant (**Supplementary Fig. 8**), supporting a broader role for the importin α5 S1B domain in STAT recognition.

**Figure 8.**
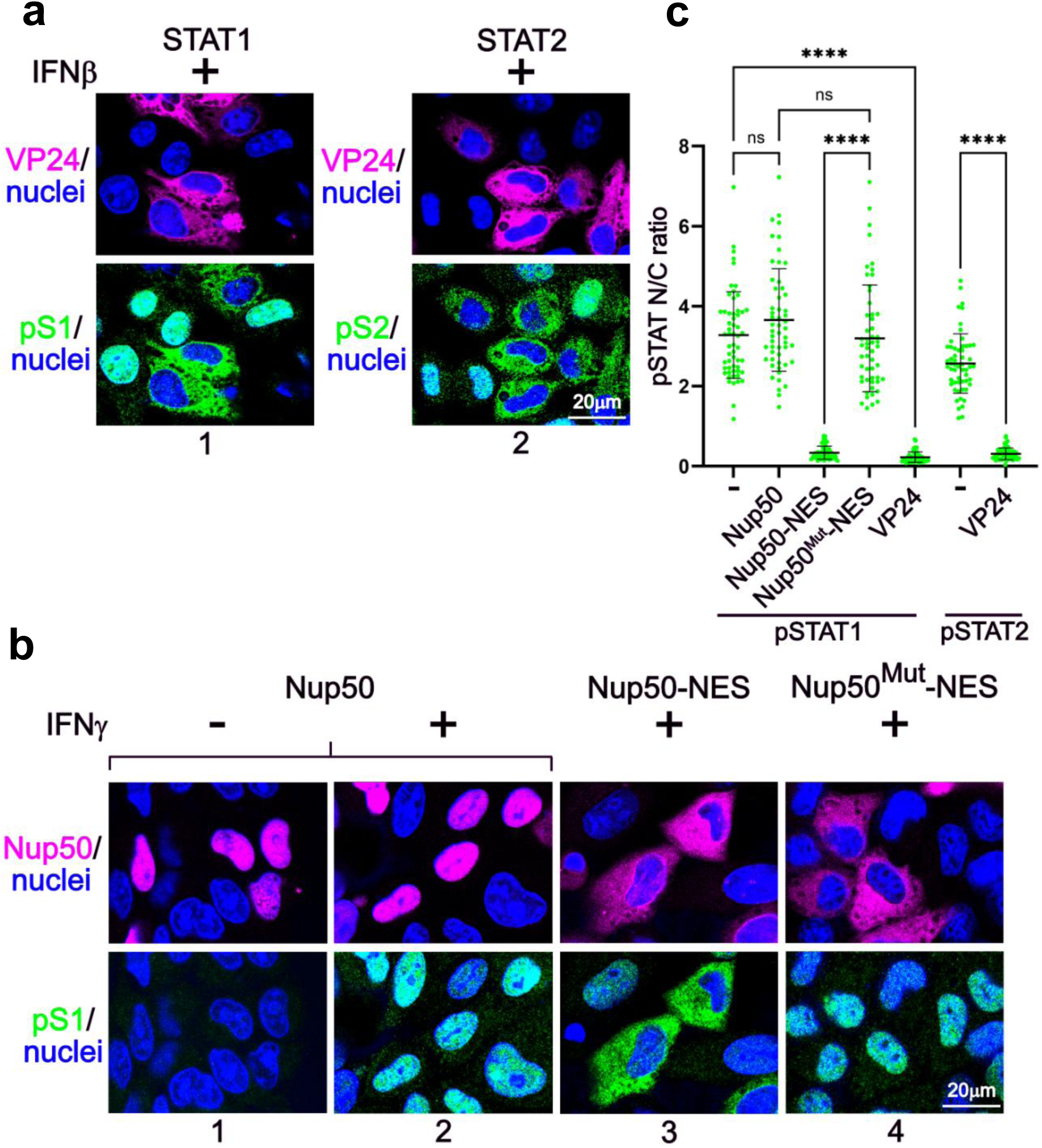
VP24 and Nup50-mediated inhibition of pSTAT1/pSTAT2 trafficking. (**a**) Confocal micrographs of HeLa cells transiently expressing VP24-mCherry fusion proteins (magenta). The distributions of VP24 fusions and endogenous Tyr701-phosphorylated STAT1 (green, row 1) or Tyr690-phosphorylated STAT2 (green, row 2) are shown after IFN-β treatment (1 h). Nuclear chromatin is shown in blue (Hoechst stain). (**b**) Cytoplasmic Nup50 inhibits IFN-γ signaling through competition with pSTAT1 for importin α5. Confocal micrographs of HeLa cells transiently expressing mCherry fusion proteins (magenta) of Nup50, NES-fused WT or mutant (Δ24–51) Nup50. The distributions of Nup50 fusions and endogenous Tyr701-phosphorylated STAT1 (green) are shown before (row 1) and after (rows 2–4) IFN-γ treatment (1 h). Nuclear chromatin is shown in blue (Hoechst stain). (**c**) Quantification of pSTAT1 and pSTAT2 nuclear accumulation (pSTAT nuclear/cytoplasmic fluorescence ratios) of cells shown in (a, b). ∗∗∗∗p <0.0001 as determined by a Kruskal–Wallis test; ns, not significant.

Activated STAT1 competes for importin α5 with VP24 in the cytoplasm and with Nup50 in the nucleus. To investigate the importance of this difference for nuclear import of pSTAT1, we engineered WT and mutant Nup50 constructs with nuclear export activity, thereby relocalizing Nup50 to the cytoplasm, and compared their effects on pSTAT1 nuclear import with those of the viral antagonist VP24 (**Fig. 8b**–**c**). Strikingly, cytoplasmic WT Nup50 potently inhibited pSTAT1 nuclear accumulation, to an extent comparable to VP24, while STAT1 tyrosine phosphorylation remained unchanged. This inhibition strictly depended on Nup50 binding to importin α5, as deletion of the α5-interacting region abolished the effect. Together, these results demonstrate that although Nup50 can compete with pSTAT1 for importin α5 in vitro and in cells, its endogenous nuclear localization renders it functionally irrelevant to pSTAT1 nuclear import under physiological conditions. Forced relocalization of Nup50 to the cytoplasm, however, converts it into a potent inhibitor of pSTAT1 nuclear import and downstream interferon signaling.

### The nuclear export factor CAS displaces pSTAT1 from importin α5

CAS (Cellular Apoptosis Susceptibility protein) functions as the export receptor for importin α, mediating its recycling from the nucleus to the cytoplasm in a Ran–GTP-dependent manner ^47^. A crystal structure of the *Saccharomyces cerevisiae* homolog of CAS (Cse1p) bound to Ran– GTP and importin α (PDB ID: 1WA5) ^48^ has revealed that CAS wraps around both Ran–GTP and the C-terminal Arm repeats 9–10 of importin α, while the IBB domain folds back to occupy the NLS-binding groove. To test whether CAS terminates pSTAT1 nuclear import, we purified recombinant CAS and performed an *in vitro* displacement assay (**Fig. 9a** and **Supplementary Fig. 5d**). As controls, we first assembled the pSTAT1–α5 complex (**Fig. 9a**, lane 5) and the ternary CAS–Ran–GTP–α5 complex (**Fig. 9a**, lane 6), which represents the importin α5 export complex. We then titrated CAS and Ran–GTP into the pSTAT1–α5 complex (**Fig. 9a**, lanes 7– 9). Sub-stoichiometric amounts of CAS and Ran–GTP only partially disassembled the pSTAT1– α5 complex (**Fig. 9a**, lane 7), while stoichiometric or excess levels of CAS and Ran–GTP completely released pSTAT1 and resulted in the CAS–Ran–GTP–α5 complex (**Fig. 9a**, lanes 8– 9). These results indicate that CAS is the likely physiological factor responsible for terminating pSTAT1 import.

**Figure 9.**
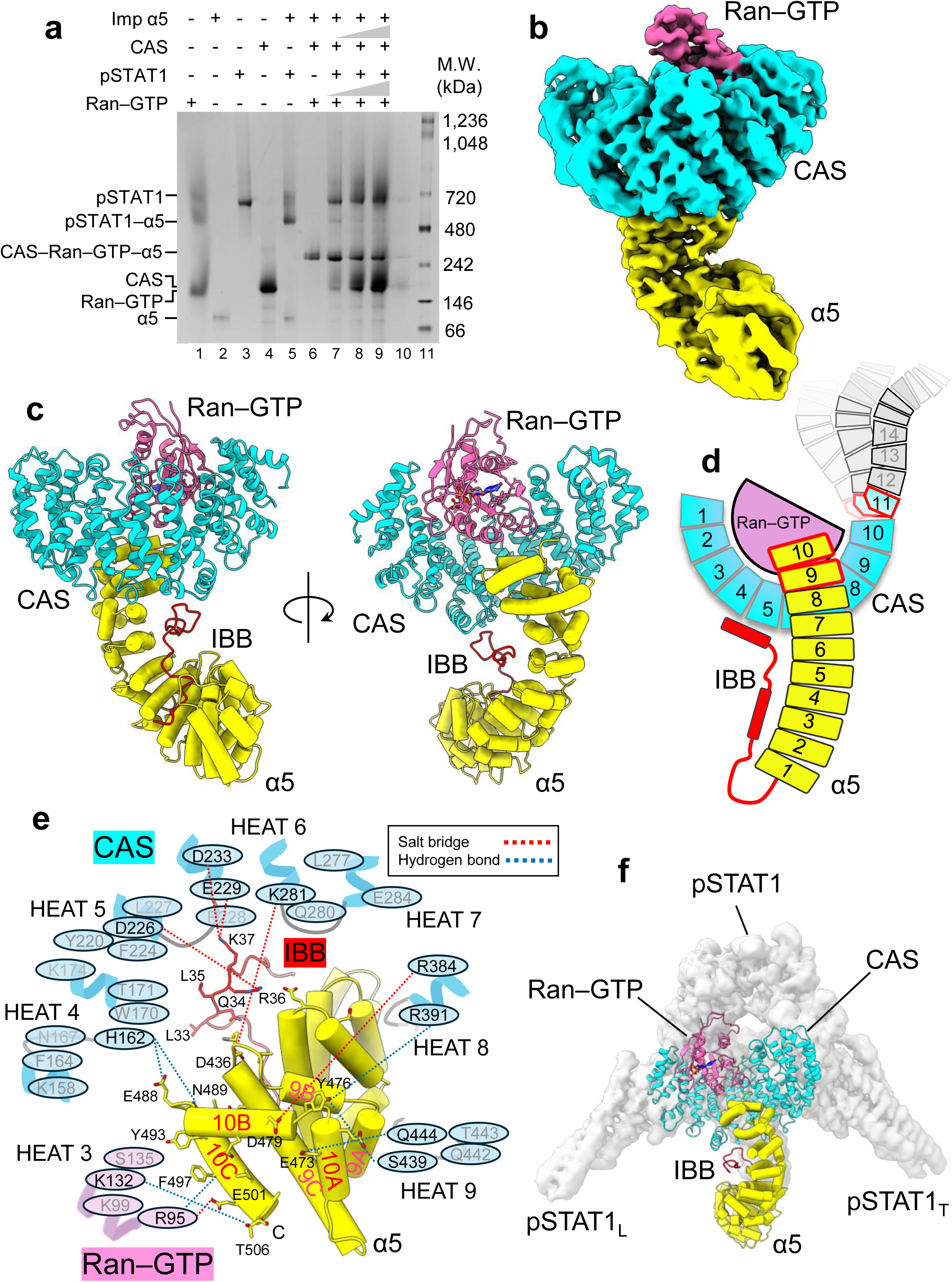
CAS and Ran–GTP drive dissociation of pSTAT1 from importin α5. (**a**) Native gel electrophoresis showing that Ran–GTP (lane 1), together with CAS (lane 4), displaces pSTAT1 (lane 3) from the pSTAT1–α5 complex (lane 5) in a concentration-dependent manner (lanes 7– 9). Titration results in formation of a trimeric CAS–Ran–GTP–α5 complex, equivalent to the control in lane 6, accompanied by excess Ran–GTP, CAS, and pSTAT1, consistent with the respective controls in lanes 1, 4, and 3. (**b**) Cryo-EM reconstruction of the CAS–Ran–GTP–α5 complex at 3.2 Å resolution. (**c**) Ribbon representation of the CAS–Ran–GTP–α5 complex, with CAS in cyan, Ran–GTP in pink, and importin α5 in yellow (IBB domain in red). The complex on the left is oriented as shown in the reconstruction in panel (**b**), while the one on the right is rotated 180° clockwise. (**d**) Cartoon representation highlighting CAS HEAT repeats involved in binding Ran–GTP and importin α5 (in cyan). HEAT repeats not visible in the reconstruction are colored gray. (**e**) Schematic of the tripartite interface between the importin α5 S1B domain (yellow), CAS HEAT repeats 3–9 (cyan), and Ran–GTP (pink). CAS and Ran–GTP residues within 4.0 Å of importin α5 are shown as ovals, whereas interacting importin α5 side chains are depicted as sticks. Dashed lines indicate salt bridges (red) and hydrogen bonds (blue). (**f**) Structural overlay of the pSTAT1–α5 complex onto the CAS–Ran–GTP–α5 complex, with pSTAT1 shown as a semitransparent surface and CAS, Ran–GTP, and importin α5 shown as ribbons.

To understand the structural basis of pSTAT1 displacement, we determined the cryo-EM structure of the CAS–Ran–GTP–α5 export complex at 3.2 Å resolution (**Supplementary Fig. 9, Fig. 9b**, and **Table 1**). The cryo-EM map enabled atomic modeling of full-length importin α5, including IBB residues 20–54, Ran–GTP, and CAS residues 1–528 (**Fig. 9c** and **Supplementary Fig. 10a**). Only CAS HEAT repeats 1–10 were well resolved, with weak density for HEAT 11 and no density for HEAT repeats 12–20, which are disordered and do not participate in importin α5 binding (**Fig. 9d**). CAS positions Ran–GTP between HEAT repeats 1–3, 6, and 9 (**Supplementary Fig. 10b**) in a baseball-in-a-glove configuration similar to that of the importin β–Ran–GTP complex ^24^. In parallel, CAS HEAT repeats 4–6 and 9–10 engage importin α5 Arm repeats 8–10 and the IBB domain (**Supplementary Fig. 10c**). Together, CAS and Ran–GTP form an extensive binding interface with importin α5 that is more extensive than that observed with pSTAT1 (**Fig. 3a**) and includes six salt bridges, seven hydrogen bonds, and 20 van der Waals contacts (≤ 4.0 Å) with importin α5 residues (**Fig. 9e** and **Supplementary Table 1**). Compared with the pSTAT1 interface, the CAS–Ran–GTP interface with importin α5 is less hydrophobic and is instead stabilized by an extensive network of polar and electrostatic interactions. These include two salt bridges and five hydrogen bonds with the S1B domain, three salt bridges with the IBB domain, and one salt bridge and two hydrogen bonds involving Ran– GTP (**Fig. 9e**).

Structural superposition of pSTAT1 and the CAS–Ran–GTP complex onto importin α5 (**Fig. 9f**) shows that CAS HEAT repeats 4–6 interact with the S1B domain similarly to the pSTAT1 DBD_L_ and ND_T,_ while HEAT repeats 9–10 mimic pSTAT1 DBD_T_. Thus, both ligands, pSTAT1 and CAS–Ran–GTP, directly compete for S1B binding and are sterically mutually exclusive (**Fig. 9f**). Consistent with the displacement assay (**Fig. 9a**), the greater number of stabilizing interactions formed by CAS–Ran–GTP (six salt bridges, seven hydrogen bonds, and 20 van der Waals) compared to pSTAT1 (four salt bridges, four hydrogen bonds, and 23 van der Waals) explains efficient cargo release. Together, these findings establish the exportin CAS as the molecular trigger for pSTAT1 release upon nuclear entry.

## DISCUSSION

Many eukaryotic signaling pathways converge on the regulated nuclear translocation of transcription factors, a process essential for coupling extracellular cues to gene expression. Despite over three decades of STAT1 biology, it remains unclear how this vital transcription factor is imported into the nucleus upon phosphorylation in the absence of a canonical NLS. Here, we define the molecular mechanism underlying STAT1 nuclear import. Our study provides a structural framework, supported by biochemical and cellular analyses, that elucidates the STAT1 nuclear import pathway, revealing how core components of the transport machinery cooperate to translocate this essential transcription factor via a non-transferable NLS.

Using cryo-EM, we found that full-length pSTAT1 has a noticeably asymmetric structure, unlike the symmetric crystal structures of DNA-bound pSTAT1 and STAT3 cores that lack NDs ^3,49,50^. Our reconstructions reveal that pSTAT1 forms an asymmetric nuclear import complex with importin α5, predominantly at a 2:1 stoichiometry at low concentrations, in which two pSTAT1 protomers engage a single importin α5 molecule. A 4:2 population is more abundant at higher concentrations and is stabilized by swapped NDs, with pSTAT1 adopting a tetrameric conformation similar to that observed upon DNA binding ^35^. As previously suggested ^32^, the NDs are critical mediators of pSTAT1 tetramerization on DNA. However, our data indicate that only a single ND is required to specifically engage importin α5, which originates from the tight protomer within the asymmetric dimer. Although all STATs harbor NDs, the dissociation constants of isolated NDs vary by more than three orders of magnitude, indicating striking differences in ND–ND interaction strength within the family ^31^. This variability supports a general model in which a single ND suffices to mediate nuclear translocation, providing a unifying mechanism for STAT nuclear import while permitting flexibility in dimerization and higher-order assembly. Consistent with this model, STAT1 bearing the double mutation Phe77Ala/Leu78Ala, which disrupts ND_T_–ND_L_ dimerization, undergoes efficient nuclear import ^45^, underscoring that the ND_T_ alone is sufficient for translocation. By contrast, deletion of the NDs abrogates importin α binding and nuclear accumulation ^20,22^. Thus, our findings highlight the importance of studying full-length macromolecules. Deletion of the NDs facilitates crystallization but eliminates asymmetry, thereby obscuring key structural features necessary to understand importin isoform specificity and the mechanistic basis of STAT1 nuclear import.

Our study elucidates the molecular basis of isoform-selective recognition of pSTAT1. Using cryo-EM, we uncovered an unexpected tripartite interface between the importin α5 S1B domain and the N-terminal and DNA-binding domains (ND_T_ and DBDs) of pSTAT1, revealing contacts unique to this isoform. Although the DBDs account for most of the interactions with importin α5 in the final pSTAT1–α5 assembly, the lack of detectable binding between importin α5 and the pSTAT1 core lacking ND_T_, together with the inability of this truncated form to support pSTAT1 nuclear import in cells ^20,22^, suggests that DBD engagement is primed by ND_T_ binding to importin α5. Our cryo-EM structures reveal that apo-pSTAT1 adopts a conformation similar to that observed in the pSTAT1–α5 complex but is slightly more open and lacks visible NDs, consistent with their intrinsic mobility in the absence of importin α5. ND_T_ engagement with importin α5 is accompanied by a subtle but potentially important closure of the pSTAT1 dimer, akin to a nutcracker motion, with the angle between the two protomers decreasing from 72.4° to 71.8°. We hypothesize that this conformational change positions the DBDs for asymmetric engagement of importin α5 Arms 9–10. This model is consistent with the inability of the pSTAT1 core and isolated NDs to support importin α5 binding when provided in trans ^20^. Rather, we propose that ND_T_ functions in cis, first recognizing Arm 10 and then promoting closure of the pSTAT1 core around the C-terminal region of importin α5, thereby stabilizing the final transport complex.

Stabilization of the ND_T_ within the pSTAT1 dimer is enabled by a structurally extended inter-repeat loop between Arm repeats 9–10 of importin α5, a feature conserved among clade 3 importin α isoforms, which promotes conformational wedging of importin α5 at the entrance to the pSTAT1 DNA-binding surface. This shape-mediated induced fit is then stabilized by two residues unique to importin α5, Lys426 (Arm 9 helix B) and Tyr476 (Arm repeats 9–10 loop), which directly contact the ND_T_. Additional importin α5 residues on the surfaces of Arm 10 helices B and C further cement the interaction with the pSTAT1 DBD_L_. Thus, isoform specificity arises from a limited set of asymmetric bonds that stabilize the pSTAT1 ND_T_ through residues absent in other importin α isoforms. The remaining interface that locks the DBDs in place likely reflects an additional induced-fit adjustment that completes the assembly of the pSTAT1–α5 complex. Notably, previous studies of importin α3 specificity for influenza A PB2 ^18^, RCC1 ^51^, and NF-κB ^52^ suggested increased conformational flexibility of this isoform relative to other importin α family members. However, recognition of these specialized cargos still depends on binding of basic NLS motifs within the Arm core, mediated primarily by polar and electrostatic interactions. By contrast, pSTAT1, to our knowledge, represents the first import cargo that completely lacks a cNLS and instead engages a distinct surface on importin α5 outside Arm repeats 2–8, suggesting the existence of additional cargos that may use a similar recognition mechanism. Thus, our findings expand the biology of importin α, establishing that its C-terminal region functions as a *bona fide* cargo-binding site. The last two Arm repeats, which together define the S1B domain, form a secondary cargo-binding surface, distinct from the major and minor NLS-binding sites (Arm repeats 2–8), which are generally occupied by cNLS cargo or the IBB domain. This finding expands the traditional view of importin α as a bipartite molecule composed of an N-terminal IBB domain and a central Arm-repeat core ^19^. Instead, importin α, at least the clade 3 isoforms, functions as a tripartite molecular platform composed of an N-terminal IBB, a central Arm core, and a C-terminal S1B domain, each contributing to distinct and precisely regulated cellular functions. Structural comparison of pSTAT1–α5 with the Ebola virus VP24–α6 complex further indicates that viral antagonists target the same C-terminal interface to disrupt host STAT1 signaling. Importin α6, a clade 3 isoform sharing >80% sequence identity with importin α5, binds VP24 at the same Arm repeats 9–10 region engaged by the pSTAT1 ND_T_ and DBDs. These findings, corroborated by biochemical and functional data for STAT2, establish the S1B domain as a critical regulatory platform at the intersection of nuclear transport and viral immune evasion. Our study provides compelling evidence that dissociation of the pSTAT1–α5–β import complex is mechanistically distinct from that of cNLS cargo. Gamma-Activated Sequence (GAS) site abundance alone cannot account for the quick nuclear accumulation of pSTAT1. Although Ran–GTP is essential for nuclear import ^23^, it is insufficient to release pSTAT1 from importin α5. This reflects the architectural segregation of the S1B domain on importin α5 (Arm repeats 9–10) from the cNLS-binding groove displaced by IBB-mediated autoinhibition (Arm repeats 2–8). CAS, not Ran–GTP alone, displaces pSTAT1 from the S1B domain upon nuclear entry, after which the trimeric CAS–Ran–GTP–α5 complex is exported and recycled ^47^. Thus, the importin α export receptor CAS, which is essential for cell viability ^47^, also plays a crucial role in human cells by terminating the pSTAT1 nuclear import and releasing the activated transcription factor into the nucleus. The coupling of cytokine-stimulated nuclear translocation of pSTAT1 with karyopherin-mediated nuclear export of importin α5 suggests an evolutionarily conserved, housekeeping mechanism for pSTAT1 nuclear trafficking that exploits established nuclear import and export pathways and the high abundance of karyopherins.

We further establish that engagement of the importin α5 C-terminus is not, in itself, sufficient to inhibit pSTAT1 nuclear import. Although VP24 and Nup50 bind a region of importin α5 that overlaps with the pSTAT1 interface, only VP24 functions as a potent antagonist, blocking IFN-induced STAT1 and STAT2 nuclear accumulation. This distinction reflects VP24’s strictly cytoplasmic localization, in contrast to Nup50’s steady-state residence at the nuclear basket. Thus, cytoplasmic localization and S1B association are both necessary conditions for inhibiting STAT1 signaling. Notably, although Nup50 efficiently displaces pSTAT1 from importin α5 in vitro, our data provide no evidence that Nup50 plays a role in STAT1 nuclear translocation in living cells ^44^. Together, these results define CAS-mediated displacement of the pSTAT1 import complex as a key mechanism terminating pSTAT1 import while ensuring efficient recycling of importin α5 for subsequent transport cycles ^53^. Whether these mechanisms of import-complex assembly and disassembly extend to additional STAT proteins remains to be determined; however, the high degree of structural conservation across the STAT family suggests that the principles identified here may be broadly applicable ^54^. In conclusion, our study defines the structural and mechanistic basis of selective, signal-dependent STAT1 nuclear import and release.

## METHODS

### Cloning, expression, and purification of recombinant proteins

The expression and purification of full-length importin α5 and ΔIBB-importin α5 (residues 66– 538) ^20^, importin β ^17^, pSTAT1 ^55^, MBP-tagged PB2 NLS (MBP-PB2-NLS) ^19^, Ran–GDP and Ran–GTP ^24^, and CAS ^53^ were carried out as previously reported, with the following modifications. The ΔIBB-importin α5 construct was cloned into pMAL-c2E and expressed in E. coli BL21-AI (Invitrogen). Cultures were induced with 0.2 mM isopropyl β-D-1-thiogalactopyranoside (IPTG) followed by 0.1% arabinose and incubated overnight at 18 °C. Harvested cells were resuspended in lysis buffer (20 mM Tris-HCl, pH 8.0, 150 mM NaCl, 3 mM β-mercaptoethanol, 1 mM phenylmethylsulfonyl fluoride (PMSF), and 0.2% Tween-20) and disrupted by sonication. Cell debris was removed by centrifugation at 25,000 × g for 30 min at 4 °C. The clarified lysate was loaded onto amylose resin (GoldBio), washed extensively with purification buffer (20 mM Tris-HCl, pH 8.0, 150 mM NaCl, 3 mM β-mercaptoethanol, 1 mM PMSF), and eluted with 10 mM maltose in lysis buffer. The eluate was dialyzed against Buffer A (20 mM Tris-HCl, pH 8.0, 50 mM NaCl, 3 mM β-mercaptoethanol, 1 mM PMSF) and further purified by anion-exchange chromatography on a HiTrap Q HP 5 mL column (Cytiva) using a linear gradient up to 1 M NaCl (Buffer B). Final polishing was achieved by size-exclusion chromatography (Superdex 200 16/600, GE Healthcare) equilibrated in gel-filtration buffer containing 20 mM Tris-HCl, pH 8.0, 150 mM NaCl, 5 mM β-mercaptoethanol, 1 mM EDTA, and 0.2 mM PMSF.

Full-length importin α5 was cloned into pGEX-4T1 and expressed in E. coli BL21-AI cells. Following induction with 0.2 mM IPTG and 0.2% L-arabinose, the cultures were grown overnight at 18 °C. Cell pellets were suspended in extraction buffer (20 mM Tris-HCl, pH 8.0, 150 mM NaCl, 3 mM β-mercaptoethanol, 1 mM PMSF, and 0.2% Tween-20) and lysed by sonication. The homogenate was clarified by centrifugation at 25,000 × g for 30 min at 4 °C, and the supernatant was applied to Glutathione Agarose resin (GoldBio). After extensive washing with buffer containing 20 mM Tris-HCl, pH 8.0, 150 mM NaCl, and 3 mM β-mercaptoethanol, the bound protein was eluted with 10 mM reduced glutathione in the same buffer. The eluted protein was dialyzed against Buffer A (20 mM Tris-HCl, pH 8.0, 50 mM NaCl, 3 mM β-mercaptoethanol, 1 mM PMSF) and further purified by anion-exchange chromatography on a HiTrap Q HP 5 mL column (Cytiva) using a NaCl gradient to 1 M. Final purification was performed by gel filtration on a Superdex 200 16/600 column (GE Healthcare) equilibrated in 20 mM Tris-HCl, pH 8.0, 150 mM NaCl, 5 mM β-mercaptoethanol, 1 mM EDTA, and 0.2 mM PMSF.

The importin α1–β heterodimer was co-expressed from a pACYCDuet-1 vector in E. coli BL21(DE3) cells. Cultures were induced with 0.5 mM IPTG and incubated for 3 h at 30 °C. Protein purification was performed as previously described ^24,56^ with the following modifications. The untagged importin β was separated from the heterodimeric complex by immobilized metal affinity chromatography (IMAC) on Nickel–NTA agarose (GoldBio) using separation buffer containing 20 mM Tris-HCl, pH 8.0, 150 mM NaCl, 250 mM MgCl_2_, and 3 mM β-mercaptoethanol. The isolated importin β fraction was further purified by size-exclusion chromatography (Superdex 200 16/600, GE Healthcare) in 20 mM Tris-HCl, pH 8.0, 150 mM NaCl, 1 mM PMSF, and 3 mM β-mercaptoethanol.

The cDNA of human STAT1α was furnished at the 3′ end with sequence encoding a 15-residue linker and the 8-residue Strep-Tag II (IBA Lifesciences), cloned into the baculovirus transfer vector pFastBac1 (Invitrogen), and expressed in baculovirus-infected Sf9 insect cells as described ^8^. Purification of the Strep-tagged protein was done according to a protocol provided by IBA Lifesciences. Unphosphorylated STAT1 was alkylated, Tyr701-phosphorylated, and purified by Heparin-affinity chromatography as described in ^8^. Tyr701-phosphorylated STAT1 was gel-filtered in phosphate-buffered saline (PBS; pH 7.4) on a Superose 12 column (GE Healthcare); peak fractions were pooled, precipitated in 50% ammonium sulfate, and the precipitate was stored at 4 °C. Before use, the protein was dissolved in PBS (pH 7.4).

MBP-PB2-NLS was cloned into a pET28a vector and expressed in E. coli NiCo21(DE3) (NEB) ^18^. Protein expression was induced by adding 0.2 mM IPTG, followed by overnight incubation at 18 °C. The harvested bacterial pellet was suspended in lysis buffer containing 20 mM Tris-HCl, pH 8.0, 150 mM NaCl, 3 mM β-mercaptoethanol, 1 mM PMSF, and 0.2% Tween-20, and lysed by sonication. After centrifugation at 25,000 × g for 30 min at 4 °C to remove insoluble material, the supernatant was applied to amylose resin (GoldBio). The column was thoroughly washed with buffer containing 20 mM Tris-HCl, pH 8.0, 150 mM NaCl, 3 mM β-mercaptoethanol, and 1 mM PMSF, and the bound protein was eluted with 10 mM maltose in the lysis buffer. The eluted fractions were pooled and dialyzed against Buffer A (20 mM Tris-HCl, pH 8.0, 50 mM NaCl, 3 mM β-mercaptoethanol, 1 mM PMSF), then subjected to anion-exchange chromatography on a HiTrap Q HP 5 mL column (Cytiva) with a linear gradient extending to 1 M NaCl (Buffer B). The final purification step consisted of gel filtration using a Superdex 200 16/600 column (GE Healthcare) equilibrated in 20 mM Tris-HCl, pH 8.0, 150 mM NaCl, 5 mM β-mercaptoethanol, 1 mM EDTA, and 0.2 mM PMSF.

Ran–GTP was purified from the Ran Q69L mutant (locked in the GTP-bound conformation) cloned into a pET28a vector and expressed in E. coli BL21-DE3 ^24^. Bacterial cells were induced with 0.25 mM IPTG and incubated at 18 °C overnight. Cells were lysed in Ran lysis buffer (50 mM KPO₄, pH 7.0, 250 mM NaCl, 2 mM MgCl_2_, 1 mM PMSF, 3 mM 2-mercaptoethanol). The lysate was applied to nickel agarose beads (GoldBio), washed with buffer containing 10 mM imidazole, and eluted in Ran elution buffer (same as lysis buffer supplemented with 150 mM imidazole). Ran–GTP was subjected to SEC on a Superdex 200 16/600 column in SEC buffer (50 mM KPO₄, pH 7.0, 250 mM NaCl, 2 mM MgCl_2_, 1 mM PMSF, 3 mM 2-mercaptoethanol). To make Ran–GDP, we expressed the full-length human Ran cloned into a pET28a-PPase vector with an N-terminal 6 × His tag in BL21-DE3 *E. coli* ^57^. The protein was expressed overnight at 18 °C in the presence of 0.5 mM IPTG. Cells were resuspended in 20 mM Tris-HCl, pH 8.0, 250 mM NaCl, 2 mM MgCl_2_, 1 mM PMSF, and 3 mM β-mercaptoethanol, lysed by sonication, and the soluble fraction was bound to low-density nickel agarose beads (GoldBio). The protein was washed with a lysis buffer containing 10 mM imidazole, then eluted with a lysis buffer (20 mM Tris-HCl, pH 8.0, 250 mM NaCl, 2 mM MgCl_2_, 1 mM PMSF, 3 mM 2-mercaptoethanol, and 150 mM imidazole). Ran was incubated with 2 mM GDP on ice for 30 min before SEC. Ran–GDP was further purified by SEC using a Superdex 200 16/600 preparative column (Cytiva) equilibrated with 20 mM Tris-HCl, pH 8.0, 250 mM NaCl, 2 mM MgCl_2_, 3 mM β-mercaptoethanol, and 1 mM PMSF.

CAS was expressed from a full-length construct cloned into the pGEX-4T1 vector, which contains an N-terminal GST tag separated by a Tobacco Etch Virus (TEV) protease cleavage site. Selection was performed via the ampicillin resistance gene. Protein expression was carried out in E. coli BL21-AI (Invitrogen) and induced by adding 0.4 mM IPTG, followed by 0.2% arabinose. Cells were harvested, resuspended in lysis buffer (20 mM Tris-HCl, pH 8.0, 150 mM NaCl, 3 mM β-mercaptoethanol, 1 mM PMSF, and 0.2% Tween-20), and lysed by sonication. The lysate was clarified by centrifugation at 25,000 × g for 30 min at 4 °C. The supernatant was incubated with Glutathione Agarose resin (GoldBio), washed extensively with buffer containing 20 mM Tris-HCl, pH 8.0, 150 mM NaCl, and 3 mM β-mercaptoethanol, and the bound GST-tagged protein was eluted with 10 mM reduced glutathione in the same buffer. The eluate was dialyzed against Buffer A (20 mM Tris-HCl, pH 8.0, 50 mM NaCl, 3 mM β-mercaptoethanol, and 1 mM PMSF) and further purified by anion-exchange chromatography on a HiTrap Q HP 5 mL column (Cytiva) with a 0–1 M NaCl gradient. Final polishing was performed by size-exclusion chromatography on a Superdex 200 16/600 column (GE Healthcare) equilibrated in 20 mM Tris-HCl, pH 8.0, 150 mM NaCl, 5 mM β-mercaptoethanol, 1 mM EDTA, and 0.2 mM PMSF. The GST tag was removed by overnight TEV protease cleavage, followed by GST resin–based separation; TEV protease was removed during the final size-exclusion chromatography step.

Importin complexes were reconstituted from purified components. For formation of the pSTAT1–α5–β import complex, 2-L bacterial cell pellets expressing importin α5 and importin β were combined and resuspended in lysis buffer containing 20 mM Tris-HCl, pH 8.0, 150 mM NaCl, 3 mM β-mercaptoethanol, 1 mM PMSF, and 0.2% Tween-20. Cells were lysed by sonication, and insoluble material was removed by centrifugation at 25,000 × g for 30 min at 4 °C. The clarified lysate was applied to Glutathione Agarose resin (GoldBio) and washed extensively with buffer containing 20 mM Tris-HCl, pH 8.0, 150 mM NaCl, 3 mM β-mercaptoethanol, and 1 mM PMSF. Importin α5, bound to the resin via its GST tag, was used to facilitate the formation of a complex with importin β. On-column cleavage of the GST tag with Precision Protease yielded the preformed importin α5–β complex, which was then combined with purified pSTAT1 to generate the trimeric pSTAT1–α5–β import complex. In parallel, the CAS–Ran–GTP–α5 complex was reconstituted by mixing purified importin α5 and CAS at equimolar concentrations, followed by the addition of a twofold molar excess of Ran–GTP.

### Mammalian cell culture, transient transfections, and CRISPR genome editing

293T (ECACC/Sigma 12022001) and HeLaS3 (ECACC 93021013) were grown in humidified incubators at 37 °C with 5% CO_2_ in Dulbecco’s Modified Essential Medium (Sigma D6429), supplemented with 1% (v/v) penicillin/streptomycin (Sigma P0781) and 10% (v/v) heat-inactivated fetal bovine serum (Sigma F9665). Gamma interferon was from Merck Millipore (#407306); human IFN-β (#11415–1) was from PBL Assay Science; staurosporine (Sigma S5921) was used at 500 nM. Where indicated for imaging experiments, HeLaS3 cells were transfected at ∼80% confluence using Lipofectamine LTX according to the manufacturer’s recommendations (Invitrogen) in 24-well plates with 0.6 μg DNA, 2 μL lipofectamine, and 2 μL PLUS reagent per well. Cells were fixed and prepared for confocal microscopy ∼15 hours later. 293T cells were CRISPR-Cas9 edited using a modification of the Synthego (Redwood

City, CA, USA) protocol “CRISPR editing of immortalized cell lines with RNPs using lipofection for 24-well plates” (2021). To direct Cas9 we utilized a CRISPR Gene Knockout Kit v2 for the human Nup50 gene (Synthego); a mixture of two single guide RNAs (sgRNA) 5’-GUUGAUAUCAUUGCUCCCAG and 5’-GCAGGUGGGAACAUUCUCCA that target the exon 5 region of the Nup50 gene as indicated in **Supplementary Fig. 6a**. Ribonucleoprotein (RNP) complexes were assembled by mixing 7.8 pmol sgRNA, 6 pmol Alt-R™ S.p. HiFi Cas9 Nuclease V3 (Integrated DNA Technologies) and 2 µL Cas9 Plus™ Reagent (Invitrogen) with 25 µL Opti-MEM™ I Reduced Serum Medium (Gibco) and incubating for 10 minutes at room temperature (RT). A transfection solution was prepared by mixing 3 μL Lipofectamine™ CRISPRMAX™ Cas9 Transfection Reagent (Invitrogen) with 25 μL Opti-MEM™ I Reduced Serum Medium (Gibco) and incubating for 5 min at RT. This solution was then mixed with the RNPs and incubated at RT for 25 min. 1.2 x 10^5^ 293T cells in 500 μL of media (DMEM containing 10% Fetal bovine serum and 1% penicillin/streptomycin) were mixed with the RNP-transfection solution and then divided equally between two wells of a 24-well plate, each containing 500 μL of prewarmed media. The cells were grown at 37 °C in 5% CO_2_ and expanded into a 6-well plate before either analysis or storage in liquid nitrogen. To increase the number of edits in the pool, the protocol was repeated on a freshly thawed stock of the edited pool and again expanded to a 6-well plate. To isolate individual clones, the pool of edited cells was resuspended to a concentration of 1 x 10^6^ mL^-1^ and passed through a Flowmi 40 µm Cell strainer (Bel-Art). Individual cells were then sorted into 96-well plates, containing 50 μL pre-conditioned media, using an Astrios EQ cell sorter (Beckman Coulter). After incubating for 1 hour at 37 °C/5% CO_2,_ a further 50 µL of fresh media was added to each well and then grown until being close to confluency. Surviving clones were again amplified up to a 6-well scale, and then genomic DNA was purified and analyzed by PCR and Sanger sequencing. Briefly, genomic DNA (gDNA) from approximately 1 x 10^6^ of WT or edited clones was extracted using the NucleoSpin Tissue, Mini kit for DNA and tissue (Macherey Nagel). The target region was amplified from approximately 20 ng gDNA using Q5^®^ Hot Start High-Fidelity DNA Polymerase (New England Biolabs), following the manufacturer’s protocol and using the primers 5’-ATTGTGCCTGGCCAAAATTT and 5’-CACTTCGCCCACCTCTGTAT. Cycling parameters for the PCR were as follows: an initial denaturation step at 98 °C for 2 min, followed by 40 cycles of 98 °C for 10 s, 66 °C for 30 s, and 72 °C for 30 s, and a final step at 72 °C for 2 min. The resulting PCR products were analyzed by agarose gel electrophoresis or purified using a NucleoSpin Gel and PCR Clean-up, Mini kit (Macherey Nagel), followed by Sanger sequencing using the primer 5’-ATTGTGCCTGGCCAAAATTTATTTTT. Whilst the WT gDNA produced a single PCR product of the expected size (493 bp), a clone was identified (designated A7C) that produced an amplicon of approximately 350 bp as indicated by agarose gel electrophoresis, but no presence of a WT-sized amplicon (data not shown). Sanger sequencing revealed this clone to contain deletions of 144 bp and 160 bp in the target region of the Nup50 gene, but no evidence of an unmodified WT allele (data not shown). To identify the effects of these deletions at the mRNA level, total RNA from approximately 3 x 10^6^ cells of either the edited clone or WT 293T cells was extracted using a RNeasy mini kit (Qiagen). cDNA was then synthesized with the SuperScript™ IV First-Strand Synthesis System (Invitrogen) according to the manufacturer’s protocol using an oligo d(T)_20_ primer and 5 µg of total RNA. A PCR was then performed as described above except using primers 5’-ACTAAGTCCTCTGAGTTCCG and 5’-GGCATCCTTTTTCTCCAGTAAAATTTTGTG and cDNA as the template. The products were analyzed by agarose gel electrophoresis or Sanger sequencing using the primer 5’-ACTAAGTCCTCTGAGTTCCG. The WT cDNA produced an amplicon of expected size (1485 bp), whereas the edited clone containing the gDNA deletions produced a single but slightly smaller product (data not shown). Sanger sequencing of the cDNA (**Supplementary Fig. 6b**) revealed that the deletions identified in the gDNA both appear to lead to a deletion of an 84 bp fragment in the mRNA, which corresponds to the whole of exon 5 of Nup50. This is predicted to result in a protein lacking amino acids 24– 51 of Nup50.

### Mammalian expression vectors

STAT1 with a C-terminal mEGFP (monomeric enhanced green fluorescent protein) was described ^58^; mutations were introduced using the Q5 site-directed mutagenesis kit (New England Biolabs). WT or the Δ24–51 mutant human Nup50 were PCR-cloned using cDNA obtained from 293T cells or the derived gene-edited cell line, respectively. PCR products coding for full-length and mutated Nup50 (1-468 aa) were ligated into vector pmCherry-N1 (Clontech) to generate fusion proteins with C-terminal red fluorescent protein (mCherry). Cytoplasm-localized WT and Δ24–51 mutant Nup50 variants were generated from the respective pNup50-mCherry vectors by inserting oligonucleotides encoding the nuclear export signal (NES) of protein kinase A inhibitor (PKI, ^34^NSNELALKLAGLDINK^49^) ^58^ into the *Age I* restriction site situated between the Nup50 and mCherry sequences. One NES sufficed for cytoplasmic localization of mutant Nup50; the WT required three. The cDNA for Ebola virus (subtype Zaire, strain H.sapiens-wt/GIN/2014/Kissidougou-C15) protein VP24 (codon-optimized) was purchased from Sinobiological (#VG40454-G), PCR-amplified, and inserted into pmCherry-C1 (Clontech). Mutagenesis and cloning results were confirmed by Sanger DNA sequencing. Molecular cloning details are available upon request.

### Confocal microscopy and image quantification

For staining endogenous proteins, cells were fixed for 15 min in ice-cold methanol and then blocked for 1 h in 20% (v/v) FBS/PBS prior to 15 h incubation at 4 °C with primary antibodies as follows: 0.2 µg mL^-1^ anti-STAT1 (#610186, Becton Dickinson), anti-phospho-Tyr701 STAT1 (1:1,000, #7649, Cell Signaling), anti-phospho-Tyr690 STAT2 (1:1,000, #88410S, Cell Signaling), anti-Nup50 (1:500; #ab137092, Abcam). After three washes in PBS, cells were incubated for 1 h at RT with 0.75 µg mL^-1^ of the appropriate species-specific secondary immunoglobulins conjugated to Alexa Fluor 488 (#A32723 or A32731, Invitrogen) or Alexa Fluor 568 dyes (#A11036, Invitrogen), in 20% (v/v) FBS/PBS. Following two further PBS washes, nuclei were counterstained with 5 µg mL^-1^ Hoechst 33258 for 5 min, and coverslips were mounted onto slides using Fluorescence Mounting Medium (Dako). Images were acquired with LSM Zeiss 880 Axio Observer confocal platform with Airyscan*FAST* detection system using Carl Zeiss Zen software, version 2. An oil-immersion 63× objective (NA 1.4) and pinhole setting 1 AU were used with laser lines at 405 nm (Hoechst dye), 488 nm (GFP), and 561 nm (mCherry). Manual image segmentation and fluorescence quantification (using integrated raw pixel intensities without background subtraction) were performed in ImageJ ^59^, as described in ^58^, for about 50 cells/experiment; experiments were done twice.

### Native and SDS-PAGE

Native gel electrophoresis was performed on NativePAGE™ 3–12% Bis-Tris Mini Protein Gels (Invitrogen, BN1001) under non-denaturing conditions. All reactions were assembled to a final volume of 15 μL using stoichiometric combinations of purified proteins. Binary complexes were formed at micromolar concentrations as follows: pSTAT1–α5 at 15 μM–5 μM and Nup50–α5 at 25 μM–5 μM. To assess competitive binding and potential displacement within the ternary complex, preformed binary complexes (either pSTAT1–α5 or Nup50–α5) were incubated with the third component for an additional 30 min at 4 °C, maintaining final concentrations of 5 μM importin α5, 15 μM pSTAT1, and 25 μM Nup50. PB2-NLS competition assays were performed similarly by titrating increasing concentrations of PB2-NLS (2.5, 5, and 20 μM) into the preassembled pSTAT1–α5 complex.

For the displacement study of the pSTAT1–α5–β import complex in the presence of Ran– GTP versus Ran–GDP, the preassembled importin α5–β complex was prepared at a final concentration of 5 μM and then assembled into a ternary complex with 15 μM pSTAT1. Displacement of the pSTAT1–α5–β ternary complex in the presence of Ran–GDP or Ran–GTP was assessed using an initial concentration of 5 μM Ran–GDP or Ran–GTP, followed by the addition of a two-fold excess of Ran–GTP to a final concentration of 10 μM. To displace the pSTAT1–α5 complex with the CAS–Ran–GTP complex, the preformed pSTAT1–α5 complex was prepared at a stoichiometric ratio of 15–5 μM, and the CAS–Ran–GTP complex was generated by mixing 5 μM CAS with 15 μM Ran–GTP. Differential titration was then performed using undersaturated (half) and saturated (two-fold) stoichiometric concentrations to achieve titration-based displacement of the pSTAT1–α5 complex. Samples were loaded onto 3–12% Bis-Tris native gels and electrophoresed in NativePAGE™ running buffer (Invitrogen) at 150 V for 90 min at 4 °C. Following electrophoresis, gels were stained with Coomassie Brilliant Blue G-250 and destained overnight in a solution containing 50% methanol and 10% acetic acid (v/v). All native gel assays were performed in triplicate.

### Western blotting and pulldown assays

The interaction between WT or mutant (Δ24–51) Nup50 and importin α5 was assessed with immunoprecipitation (IP) experiments using human cell line 293T or its CRISPR-edited derivative (clone A7C) expressing mutated Nup50. 1 x 10^6^ 293T cells per well of a 6-well plate were transfected (1 μg plasmid/well) with human FLAG-tagged importin α5 encoding mammalian expression plasmid ^42^ or pmGFP-N1 control plasmid ^58^ using Lipofectamine LTX according to the manufacturer’s recommendations (Invitrogen). One to two days post transfection, where indicated cells were treated for 1 hour with 50 U mL⁻¹ human IFN-γ, enzymatically dislodged (trypsin), washed in 1 mL ice-cold PBS, resuspended in 1 mL per well ice-cold PBS-T (PBS containing 0.5% Triton X-100 (Sigma T8787)) and sonicated in an ice bath using a high intensity ultrasonic processor (Sonics & Materials, Inc. Model VC50T) fitted with a 3 mm microtip for 3 cycles of 10 s at 6.25 Watts and 20 s cooling. The homogenate was centrifuged at 15,000 x g for 10 minutes, and the supernatant was added to ∼10 µL of equilibrated anti-FLAG M2 magnetic resin, which was incubated at 4 °C on a rotating shaker for 1 hour. Using a Magrack 6 (GE Healthcare 28948964) magnetic separation rack, the resin was washed four times with 1 mL of PBS-T, then residual buffer was removed, and the resin was resuspended in 30 µL of 2x SDS sample buffer before heating at 95 °C for 5 min. The whole 30 µL immunoprecipitated material (or whole-cell extract) was resolved by 10% SDS-polyacrylamide gel electrophoresis and then transferred to a nitrocellulose membrane (Amersham Protran 0.45 μm 10600002) using a Trans-Blot SD Semi-Dry Transfer Cell (BioRad) at a constant 16 Volts for 90 min. Membranes were stained with Ponceau S for imaging, fully destained in TBS containing 0.05% Tween 20 (TBS-T), and blocked in TBS-T containing 4% dehydrated milk protein (Marvel) for 1 hour at room temperature. Blots shown in Supplementary Figs. 4, 7 and 8 were probed overnight (∼16 hours) at 4 °C with anti-FLAG M2 (1:1,000; Sigma F3165), anti-human Nup50 (1:1,000; Abcam ab137092), anti-human STAT1 (1:4,000, Santa Cruz sc-345), anti-Tyr701-phosphorylated STAT1 (1:1,000, Cell Signaling 9171), anti-human STAT2 (1:1,000, Atlas antibody Merck HPAO123458), anti-Tyr690-phosphorylated STAT2 (1:1000, 88410S, Cell Signaling), or anti-β-actin (1:8,000, Sigma, A5441) primary antibodies in TBS-T, washed and probed in this buffer with IRDye 680LT Goat anti-Mouse IgG (H+L) (1:5,000; LI-COR 926-68020) and IRDye 800CW Goat anti-Rabbit IgG (H+L) (1:5,000; LI-COR 925-32211) secondary antibodies. After final washes in TBS-T, blots were imaged on an Odyssey CLx Imager (LI-COR). Anti-FLAG M2 magnetic resin (Sigma M8823) was equilibrated as follows. 60 µL of a 50% slurry of M2 magnetic affinity resin was washed with 5 mL of PBS (Sigma D8537) containing 0.1% BSA fraction V (Roche 8076.2), then with 5 mL of PBS-T. The resin was resuspended in 6 mL PBS-T, and beads equivalent to 1 mL of this were used per IP reaction. All steps were performed at 0 to 4°C.

The interactions between pSTAT1/pSTAT2 and importin α5 (WT or mutant Tyr476Ala) were similarly assessed with IP experiments using 293T cells. Cells were transfected with FLAG-tagged importin α5 as described above; where indicated, cells were treated for 1 hour with 50 U mL⁻¹ human IFN-γ or 500 U/ml IFN-β, followed by cell lysis, IP using Anti-FLAG M2 magnetic resin and Western blotting as described above.

The native gel showing the association of PB2-NLS with pSTAT1 and importin α5 was transferred onto a 0.45 μm PVDF membrane (Immobilon PVDF, IPVH00010) using a Mini Trans-Blot Cell (Bio-Rad) at a constant 30 V overnight in transfer buffer (25 mM Tris-HCl, pH 8.0, 192 mM Glycine, 20% Methanol, 0.05% SDS). Membranes were stained with Ponceau S to confirm protein transfer, imaged, fully destained in TBS-T, and blocked for 1 hour at room temperature in TBS-T supplemented with 4% (w/v) non-fat dry milk (Marvel). Blots were incubated overnight (∼16 hours) at 4 °C with mouse anti-His primary antibody (1:1,000 dilution; ABclonal, AE003), washed with TBS-T, and subsequently incubated with horseradish peroxidase (HRP)-conjugated anti-mouse secondary antibody (1:2,500 dilution; Promega, W402B). Protein bands were visualized by chemiluminescence using a luminol-based detection kit (Thermo Scientific SuperSignal™ West Pico PLUS Chemiluminescent Substrate; 34577), and signals were captured with a Bio-Rad ChemiDoc MP imaging system.

### Vitrification and data collection

To vitrify the pSTAT1–α5 complex at low concentration (LC), purified pSTAT1 and importin α5 were mixed at a 2:1 molar ratio, and a 3 μL aliquot at a final concentration of 1.0 mg mL⁻¹ was immediately applied to a gold UltrAufoil R 1.2/1.3, 300 mesh grid (EMS) that had been glow-discharged for 30 s at 15 mA using an easiGlow system (PELCO). For the high-concentration (HC) pSTAT1–α5 complex, a 3 μL aliquot of gel filtration-purified pSTAT1–α5 concentrated to 1.5 mg mL⁻¹ was applied to a glow-discharged gold UltrAufoil R 1.2/1.3, 300 mesh grid (EMS). Grids were blotted for 5 s at a blot force of 5 and vitrified in liquid ethane cooled with liquid nitrogen using a Vitrobot Mark IV (FEI). For the CAS–Ran–GTP–α5 complex, 3 μL of gel filtration-purified sample at 1.5 mg mL⁻¹ was applied to a glow-discharged UltrAufoil R 1.2/1.3, 300 mesh grid (EMS) and vitrified as described above. All grids were pre-screened in-house on a 200 kV Glacios 2 transmission electron microscope equipped with a Falcon 4i direct electron detector. Data collection on the Falcon 4i detector was performed using EPU (v3.10) software ^60^ in accurate positioning mode.

For the (LC) pSTAT1–α5 complex, we collected high-resolution data on a 300 kV Titan Krios transmission electron microscope equipped with a Falcon 4i direct electron detector at the National Center for CryoEM Access and Training (NCCAT). A total of 12,613 movies were recorded in super-resolution mode using a 20 eV energy-filter slit at a calibrated pixel size of 0.717 Å (165,000 × magnification). The nominal total exposure was 50 e⁻ Å⁻² fractionated over 40 frames with a defocus range of −0.65 to −2.5 μm. For the (HC) pSTAT1–α5 complex, high-resolution data were collected on a 300 kV Titan Krios transmission electron microscope equipped with a field-emission gun and a Gatan K3 BioQuantum direct electron detector at the Stanford–SLAC Cryo-EM Center (S2C2). A total of 19,493 movies were recorded in super-resolution mode using a 20 eV energy-filter slit at a calibrated pixel size of 0.86 Å (105,000 × magnification). The nominal total exposure was 50 e⁻ Å⁻² fractionated over 40 frames with a defocus range of −0.8 to −2.5 μm. For the CAS–Ran–GTP–α5 complex, an initial dataset of 20,004 movies was collected in-house on a 200 kV Glacios 2 microscope equipped with a Falcon 4i detector, with a nominal total exposure of 50 e⁻ Å⁻² distributed over 36 frames. A second, higher-resolution dataset of the CAS–Ran–GTP–α5 complex was collected on a Titan Krios transmission electron microscope equipped with a Gatan K3 direct electron detector and an energy filter at the National Cryo-Electron Microscopy Facility (NCEF), National Cancer Institute. A total of 21,503 micrographs were acquired at a calibrated pixel size of 0.855 Å (105,000 × magnification), with a nominal total exposure of 50 e⁻ Å⁻² fractionated over 40 frames and a defocus range of −0.8 to −2.0 μm. Data were collected using SerialEM (v4.2.10) ^61^ at S2C2 and Leginon ^62^ at NCEF. All data collection parameters are summarized in **Table 1**.

### Cryo-EM single-particle analysis

The image-processing workflow for (LC) pSTAT1–α5, summarized in **Supplementary Fig. 1**, yielded the 2:1 pSTAT1–α5 complex. All processing was performed in cryoSPARC v5.0.7 ^63^. An initial pool of 12,613 movies underwent gain-normalized patch motion correction and patch contrast transfer function (CTF) estimation. A set of 4,825 pSTAT1–α5 (LC) micrographs was retained following exposure curation using a CTF fit cutoff of 5 Å. From these micrographs, unbiased blob picking, using particle diameters of 250 and 100 Å, generated a stack of 1,615,518 particles. Micrographs affected by ethane and ice contamination or other grid artifacts were excluded using the Micrograph Junk Detector, yielding 743,391 particles. Particles were extracted in 800-pixel boxes with ½-Fourier cropping, producing a working stack of 737,783 particles, which was subjected to 2D classification with minimal cleaning to yield a stack of 717,845 particles. Visually distinct 2D class averages were isolated into independent particle stacks and used to generate initial ab initio volumes, followed by non-uniform refinement to produce references for unbiased heterogeneous refinement. Multi-class heterogeneous refinement was performed on 717,845 particles to segregate the 2:1 pSTAT1–α5 complex from contaminating, noisy, and heterogeneous species, including apo-pSTAT1 dimers and tetramers, which showed excessive heterogeneity and poor particle distribution and were not pursued further. The dedicated class of the 2:1 complex was further used for template generation and to train a Topaz picking model ^64^. A particle stack generated using the Blob, Topaz, and template-based workflows was pooled, yielding a total of 2,704,667 particles across the entire dataset. Following duplicate particle removal, several rounds of heterogeneous refinement were performed on this particle stack using the previously generated 3D volumes to obtain a set of 495,055 particles. α5-focused 3D classification was further used to generate a clean, full-resolution stack of 260,316 particles. Using non-uniform refinement with per-particle defocus refinement, 255,976 particles converged at 3.8 Å resolution based on the gold-standard FSC (0.143 criterion). The particles then underwent iterative rounds of local and global CTF correction, followed by reference-based motion correction, yielding a final stack of 246,208 particles with an overall resolution of 3.6 Å (GSFSC). The final refined map was used for local-resolution estimation and visualization.

The image-processing workflow for the (HC) pSTAT1–α5 dataset (**Supplementary Fig. 2**) yielded the 4:2 pSTAT1–α5 complex and an apo-pSTAT1 dimer. Movies were subjected to gain-normalized patch motion correction and patch CTF estimation, and all subsequent processing was performed in cryoSPARC v4.2.1 or v5.0.2 ^63^. We selected an initial 1,000 micrographs from a pool of 19,493 micrographs to generate reference templates and a preliminary reconstruction. Particles were picked using a blob picker, extracted in a small box (360 px), and cleaned by 2D classification; particles corresponding to well-resolved 2D averages were used to generate templates for template-based picking on the same subset. Template-picked particles were re-extracted (360 px) and classified by 2D classification, and selected particles were used for ab initio reconstruction, yielding a 6 Å preliminary map from 26,210 particles. This map was refined using non-uniform refinement and further cleaned with heterogeneous refinement against several noise classes, producing a preliminary reconstruction aligned to the STAT1 dimer core. Two masks were generated corresponding to a “tight” and a “loose” conformation of the STAT1 N-terminal domain (ND) for use in subsequent focused classification. Templates from this pilot dataset were then used to pick particles across the full micrograph set, yielding 5.6 million particles after extraction (360 px). Following 2D classification, particles containing a clear STAT1 dimer core were retained (1.5 million particles) and refined by non-uniform refinement on 2x-downsampled particles (3.5 Å at Nyquist). To resolve heterogeneity in importin α5 binding at the STAT1 N-terminal domain, focused 3D classification was performed independently using the ND_T_ and ND_L_ masks, resolving three populations: α5 stably bound at the loose ND conformation, α5 weakly bound at the loose ND conformation, and particles lacking density for α5/ND. The particle stack was re-extracted at full resolution, and heterogeneous refinement was performed using these three populations together with three noise classes, yielding approximately 1.1 million particles distributed among the three classes. Particles were re-extracted in a larger box (640 px) to accommodate the full 4:2 assembly, and additional focused 3D classification on the ND masks and density corresponding to a second STAT1 protomer resolved <u>five</u> distinct populations. Heterogeneous refinement using these five populations together with three noise classes yielded: a first stable 4:2 conformation (Conf 1, 199,271 particles), a second 4:2 conformation (Conf 2, 167,940 particles), a mixed population of 2:1 complex and loosely tethered 4:2 assembly (187,912 particles) (**Supplementary Fig. 3**), an apo-pSTAT1 dimer lacking importin α5 (206,115 particles), and an unstable/heterogeneous 2:1 population (185,843 particles). The apo-pSTAT1 dimer and the first stable 4:2 conformation were carried forward for final refinement. Each stack was subjected to iterative rounds of non-uniform refinement together with local and global CTF refinement. The final stable 4:2 conformation 1 map was reconstructed from 146,431 particles with C2 symmetry, at a resolution of 3.6 Å; the final apo-pSTAT1 dimer map was reconstructed from 196,697 particles with C2 symmetry, at a resolution of 3.3 Å.

To reconstruct the CAS–Ran–GTP–α5 complex (**Supplementary Fig. 9**), single-particle analysis was conducted using cryoSPARC v5.0.2 ^63^ on the 21,503 micrographs collected at NCEF. After patch-based motion correction and CTF estimation, micrographs were curated to a subset of 17,869 based on CTF fit resolution, relative ice thickness, and total full-frame motion distance. Extensive 2D classification was used to sort particles identified by blob picking from this subset of micrographs. We observed a disproportionate number of 2D classes with similar orientations. To enrich for a more diverse range of 2D views, we identified rare classes and subjected these particles to Topaz-based picking ^64^. After initial attempts to reconstruct the complex, we suspected that heterogeneity was limiting resolution and causing maps to appear heavily anisotropic. In order to identify a relatively stable conformation of the complex, we sought to first align the particles to CAS–Ran–GTP without α5. We used the molmap command in ChimeraX ^65^ to generate a simulated 3.5 Å map from a model based on a high-confidence AlphaFold3 prediction of the CAS–Ran–GTP complex, rigid-body fitted into the anisotropic map. Well-resolved classes from 2D classification were refined by non-uniform refinement using the simulated CAS–Ran–GTP map as the initial reference. The CAS–Ran–GTP resolved well, and inspection at low contour revealed poorly resolved density indicative of importin α5, giving us confidence that we were not introducing input bias, as α5 was absent from the simulated input map. As expected, only a few helices at the C-terminus of importin α5 were resolvable at the interface between CAS and importin α5, suggesting heterogeneity in the N-terminus of α5. We performed 3D classification using a focus mask around importin α5 to isolate a stable conformation of the complex. Reconstruction of the three best classes resulted in a 3.7 Å map. Homogeneous *ab initio* refinement and subsequent local refinement of this single class confirmed an almost identical complex conformation at 3.8 Å, again indicating that we were not over-biasing the density with our input map. Using a larger particle stack of ∼7 million particles, we then ran a heterogeneous refinement job using the 3.7 Å map, seven anisotropic complex maps, and four junk classes generated from a single round of *ab initio* with an initial low-pass resolution of 5 Å. This yielded a well-resolved isotropic stack of ∼800,000 particles, which, after reference-based motion correction and iterative rounds of local refinement, local CTF refinement, and 3D classification to clean up the particle stack, led to a final map of 286,636 particles with an FSC of 3.19 Å.

### Model building, refinement, and structure analysis

All cryo-EM maps generated in this study were visualized using ChimeraX ^65^ and Coot ^66^. Atomic models were built using available PDBs, AlphaFold3 ^67^ models, ISOLDE ^68^, or *de novo* using Coot ^66^. Atomic models were refined through several rounds of rigid-body, real-space, and B-factor refinement with *phenix.real_space_refinement* ^69^. Model refinement and validation were performed in Phenix, and model quality was assessed using MolProbity (v4.5.2) ^70^. Detailed statistics on data collection, refinement, and validation are provided in **Table 1**. Overall, we refined and deposited atomic coordinates for four structural models. (**i**) The asymmetric model of the pSTAT1–α5 2:1 assembly from the (LC) dataset was refined to a map-to-model correlation coefficient of 0.83 (CC_mask), calculated within a mask surrounding the atomic model, and 0.88 (CC_box), calculated over the entire map volume including solvent and empty regions, at 3.6 Å resolution (**Table 1**). (**ii**) The pSTAT1–α5 4:2 assembly from the (HC) dataset was real-space refined against the 3.6 Å map, yielding a map-to-model of 0.82 (CC-mask) and 0.91 (CC-box) (**Table 1**), after applying 2-fold rotational symmetry (C2). (**iii**) The apo-pSTAT1 from the (HC) dataset was real-space refined against the 3.3 Å map, yielding a map-to-model of 0.85 (CC-mask) and 0.92 (CC-box) (**Table 1**). (**iv**) The trimeric CAS–Ran–GTP–α5 complex was real-space refined against the 3.2 Å map, yielding map-to-model values of 0.73 (CC-mask) and 0.81 (CC-box) (**Table 1**).

All images of structures and cryo-EM maps were generated using ChimeraX ^65^ and PyMOL ^71^. RMSD between superimposed PDBs, structural comparisons, and domain motion analysis were done using the DynDom server ^72^ and GESAMT from the CCP4i2 software suite ^73^. Binding interfaces were analyzed using PISA ^74^ and PDBsum ^75^. AlphaFold3 ^67^ was used to predict an initial model of human CAS. Cartoon models were generated in part using BioRender.com, which is licensed at UAB.

### Quantification and Statistical Analysis

Statistical analyses of immunofluorescence results were performed in GraphPad Prism Software Version 9.3.0. D’Agostino-Pearson’s omnibus K2 test was used to assess the normality and log-normality of individual variables. The Kruskal–Wallis test, in conjunction with Dunn’s test to correct for multiple comparisons, was used for hypothesis testing. The ROUT method with Q = 1% was utilized to identify outliers. Significance is designated as ∗*p* < 0.05; ∗∗*p* < 0.01; ∗∗∗*p* < 0.001; ∗∗∗∗*p* < 0.0001. To estimate the resolution of all cryo-EM reconstructions in this study from Fourier Shell Correlation (FSC) curves, the FSC cut-off criterion of 0.143 ^76^ was used. Model-to-map agreement was evaluated using FSC (0.5 and 0.143) criteria and map–model correlation coefficients (CC_mask), calculated within a mask surrounding the atomic model, and (CC_box), calculated over the entire map volume, including solvent and empty regions. Local resolution was estimated using cryoSPARC v4.2.1 or v5.0.2 ^63^. No statistical methods were used to pre-determine sample size. The experiments were not randomized. The investigators were not blinded to allocation during the experiments or during outcome assessment.

## DATA AVAILABILITY

The atomic coordinates generated in this study have been deposited in the Protein Data Bank database under accession codes 38YK {https://doi.org/10.2210/pdb38YK/pdb} for the 2:1 pSTAT1–α5 complex; 11JG {https://doi.org/10.2210/pdb11JG/pdb} for the 4:2 pSTAT1–α5 complex; 38HT {https://doi.org/10.2210/pdb38HT/pdb} for the apo-pSTAT1 dimer; 11PH {https://doi.org/10.2210/pdb11PH/pdb} for the CAS:Ran–GTP:α5 complex. The cryo-EM density maps generated in this study have been deposited in the Electron Microscopy Data Bank under accession codes

EMD-79237 {https://www.emdataresource.org/EMD-79237}(2:1 pSTAT1–α5 reconstruction), EMD-75737 {https://www.emdataresource.org/EMD-75737}(4:2 pSTAT1–α5 reconstruction), EMD-78826 {https://www.emdataresource.org/EMD-78826}(apo-pSTAT1 reconstruction), EMD-75922 {https://www.emdataresource.org/EMD-75922}(CAS:Ran-GTP:α5 reconstruction).

Previously published PDB codes referred to in this paper include:

PDB 1BF5 {http://doi.org/10.2210/pdb1BF5/pdb} (dimeric pSTAT1:DNA complex), PDB 8YYV {http://doi.org/10.2210/pdb8YYV/pdb} (dimeric pSTAT1:DNA complex), PDB 8YYU {http://doi.org/10.2210/pdb8YYU/pdb} (tetrameric pSTAT1:DNA complex) PDB 1WA5 {http://doi.org/10.2210/pdb1WA5/pdb} (Cse1:Kap60:Ran–GTP complex). PDB 3TJ3 {http://doi.org/10.2210/pdb3TJ3/pdb} (Importin α5:Nup50 complex). PDB 4U2X {http://doi.org/10.2210/pdb4U2X/pdb} (Ebola virus VP24:α5 complex).

The minimum datasets (e.g., motion-corrected micrographs) required to interpret and verify all four cryo-EM reconstructions presented in this paper can be downloaded from the University of Alabama at Birmingham (UAB) Cheaha Supercomputer (https://www.uab.edu/it/home/research-computing/cheaha) upon request to the authors.

## ACKNOWLEDGMENTS

We thank Drs. Christopher Basler, Icahn School of Medicine at Mount Sinai, New York, for providing the FLAG-tagged human importin α5 expression plasmid, and Drs. Siegfried Musser, Texas A&M University, and Roderick Lim, Biozentrum, Basel, for the CAS expression plasmid. We are grateful to Dr. Chun-Feng David Hou and Ms. Fenglin Li for help with vitrifying pSTAT1– α5 grids; to Dr. James Kizziah at UAB for assisting with in-house cryo-EM data collection; to Drs. Robert Markus and Daniel Scott at UoN for imaging support and advice on CRISPR editing, respectively. Electron microscopy was conducted at the UAB Cryo-EM Facility (RRID:SCR_025450), supported by the Institutional Research Core Program and O’Neal Comprehensive Cancer Center (NIH grant P30 CA013148), with additional funding from NIH grant S10 OD024978. We also thank the School of Life Sciences Flow Cytometry Facility, and federal cryo-EM facilities for access to 300 kV Krios microscopes and assistance with data collection: the National Cancer Institute’s National Cryo-EM Facility at the Frederick National Laboratory for Cancer Research (contract 75N91019D00024); the National Center for CryoEM Access and Training (NCCAT) and the Simons Electron Microscopy Center (supported by the NIH U24 GM129539, the Simons Foundation SF349247, and NY State Assembly); and the Stanford-SLAC Cryo-EM Center (S2C2), supported by NIH U24 GM129541.

## AUTHOR CONTRIBUTION STATEMENT

U.V. and G.C. conceived the study. O.W., L.H., P.K., R.Y., J.C., and L.K.R. purified all proteins. P.K. and R.Y. performed biochemical experiments. P.K., R.Y., and S.S.S. vitrified and collected micrographs. P.K. and N.F.B. solved cryo-EM reconstructions. P.K. and G.C. built, refined, and analyzed all structural models. J.C., O.W., A.B., and U.V. carried out all cellular studies. P.K., U.V., and G.C. wrote the manuscript with feedback from all other authors.

