## Supplemental file for "Cytokine-induced nuclear translocation of STAT1 via a non-transferable NLS"

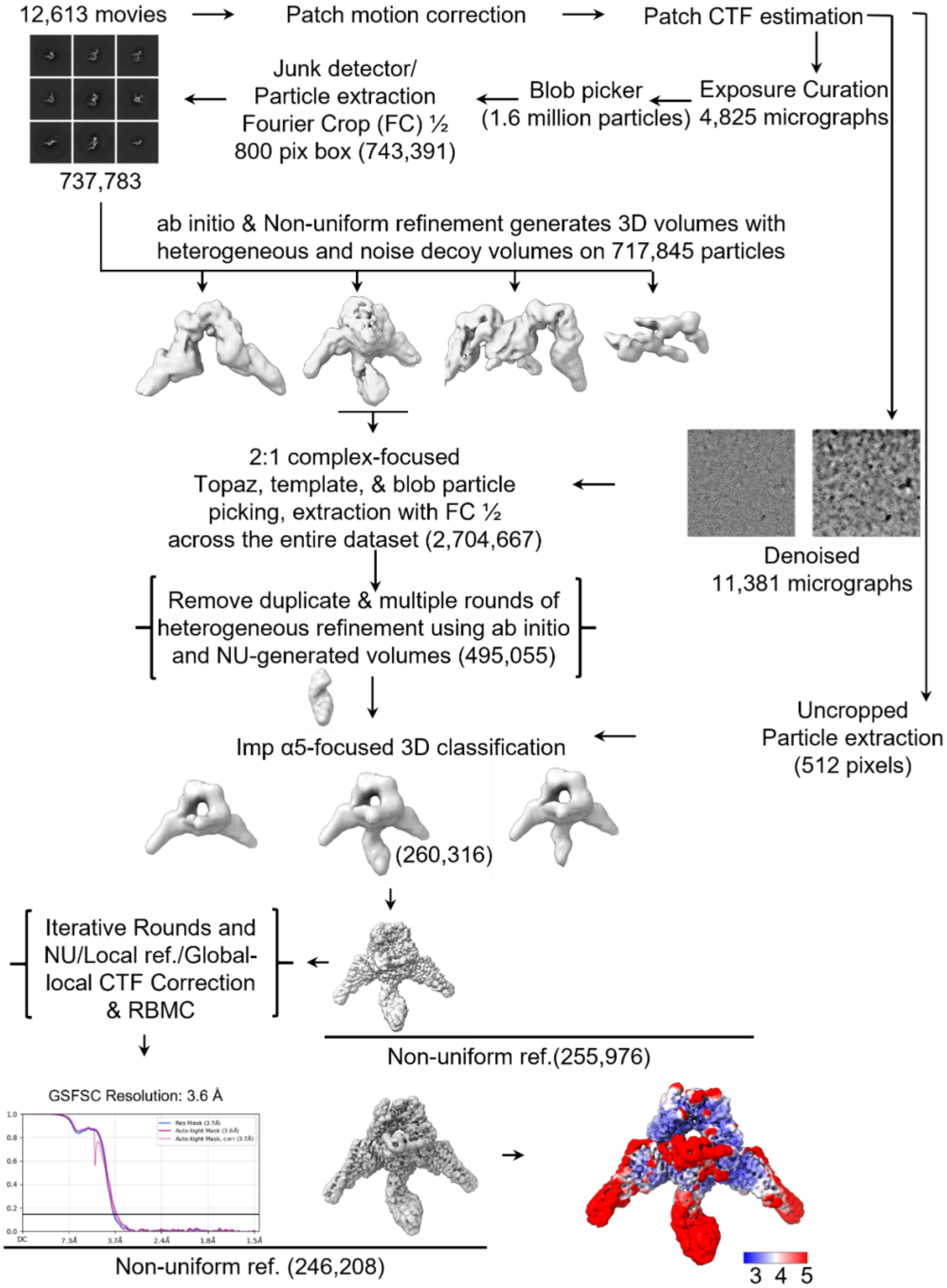

**Supplementary Fig. 1. Workflow of cryo-EM single-particle analysis for the low-concentration (LC) pSTAT1–α5 dataset.** The Fourier shell correlation (FSC) curve for the final reconstruction is shown for the 2:1 pSTAT1–α5 complex. The final reconstruction reached a global resolution of 3.61 Å at the FSC = 0.143 criterion using a cryoSPARC-automatically generated tight mask, 3.67 Å using the resolution mask, and 3.72 Å using the refinement mask employed during multi-cycle convergence.

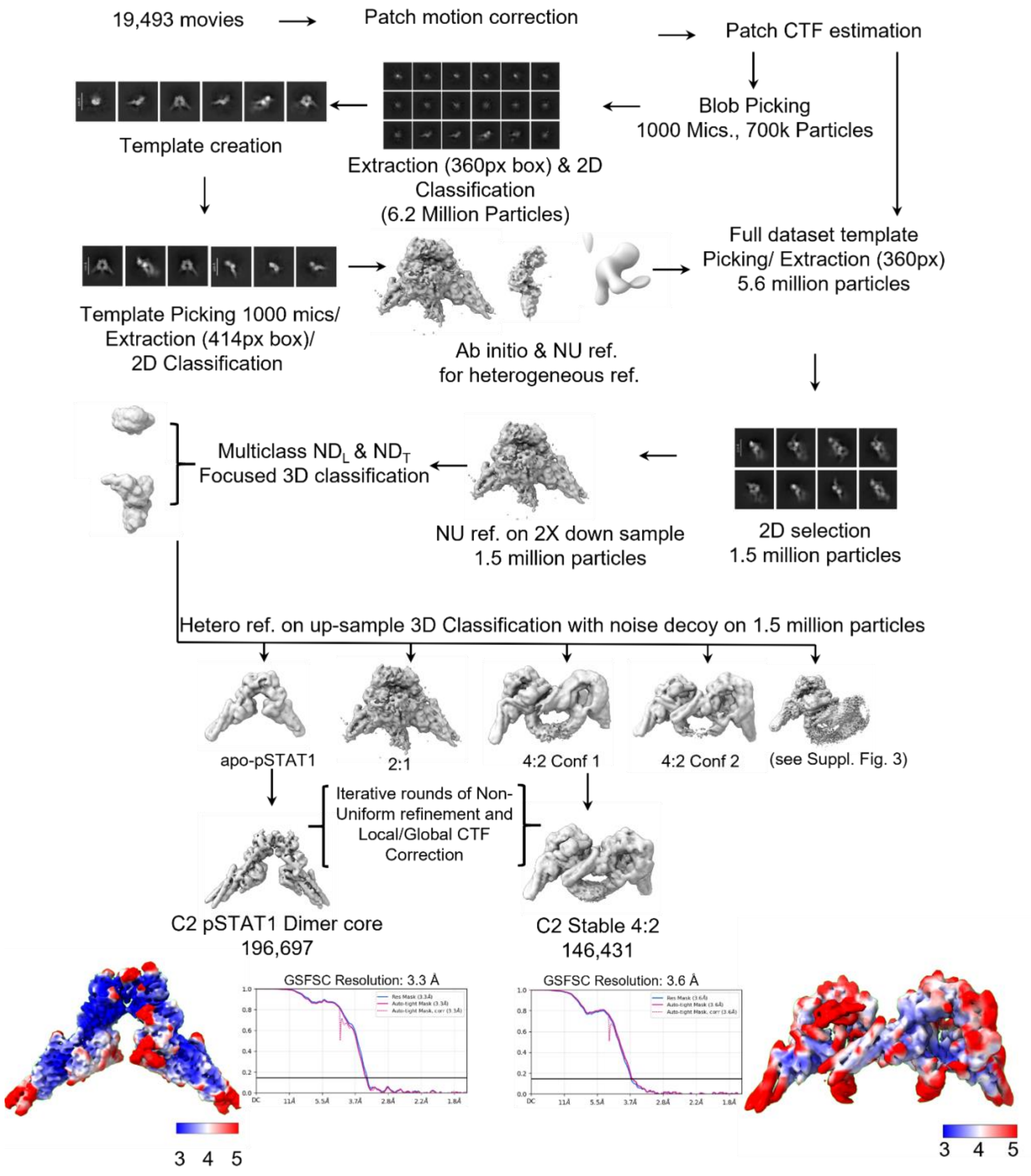

**Supplementary Fig. 2. Workflow of cryo-EM single-particle analysis for the high-concentration (HC) pSTAT1–α5 dataset.** Fourier shell correlation (FSC) curves for the final reconstructions are shown for the apo-pSTAT1 (left) and the 4:2 pSTAT1–α5 complex (right). We estimated resolution using the FSC = 0.143 criterion, yielding overall resolutions of 3.3 Å and 3.6 Å, respectively.

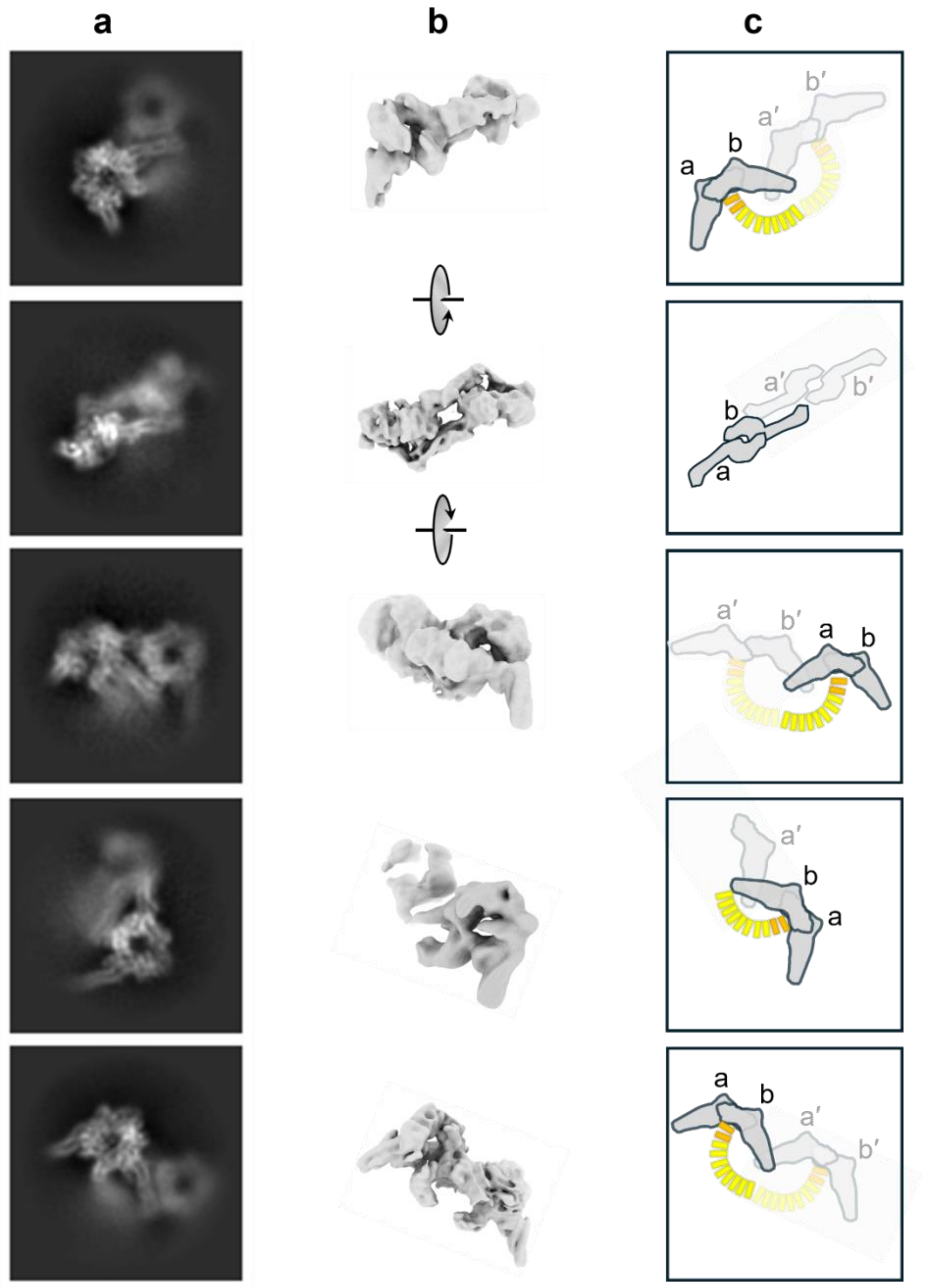

**Supplementary Fig. 3. Imperfect assembly of the tetrameric pSTAT1–α5 complex identified in the (HC) pSTAT1–α5 dataset.** Representative 2D class averages (**a**) and corresponding 3D reconstructions (**b**) of the 4:2 pSTAT1–α5 complex reveal slippage of the secondary pSTAT1 dimer (protomers a', b') relative to the primary dimer (protomers a, b) (see **Fig. 2e**). Panel (**c**) shows a schematic of the asymmetric tetrameric populations, with the primary dimer in dark gray and the slipping dimer in semi-transparent gray. Importin α5 is shown in yellow, with Arms 9–10 highlighted in orange.

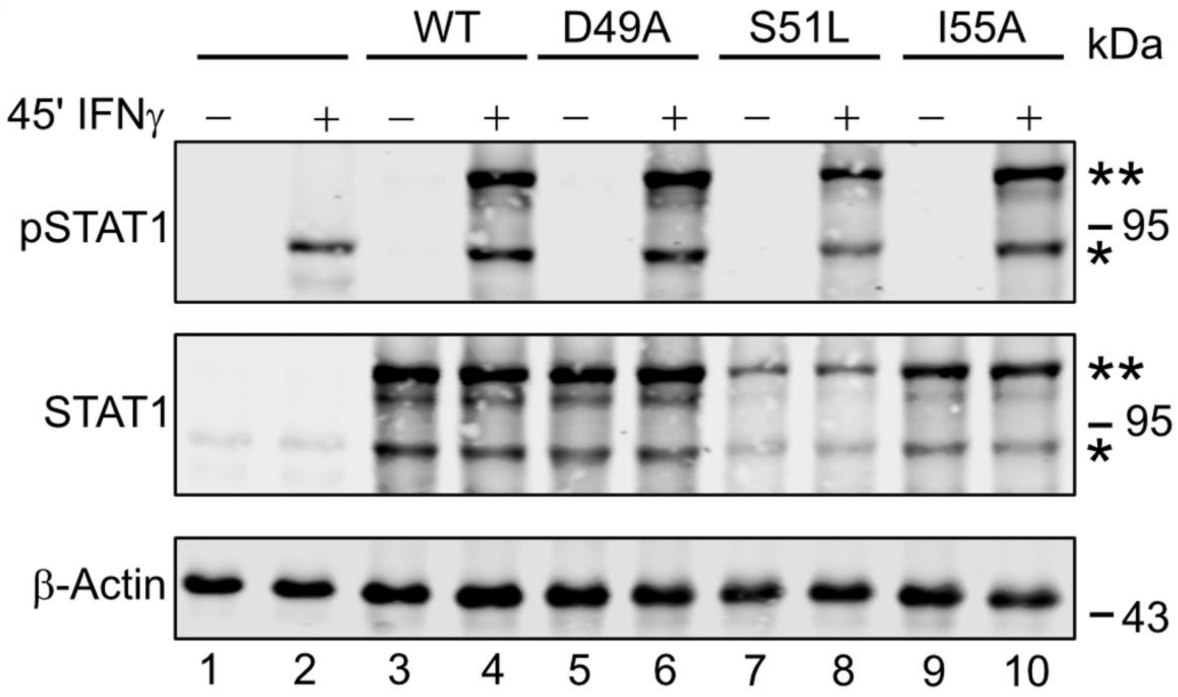

**Supplementary Fig. 4. IFN-γ-induced Tyr701-phosphorylation of STAT1 ND mutants.** Whole-cell extracts from untreated and IFN-γ-treated 293T cells (lanes 1–2) and 293T cells transfected with STAT1-GFP wild-type (lanes 3–4) or the indicated mutants (lanes 5–10) were Western-blotted and concurrently probed with pSTAT1 (top panel), STAT1 (middle panel), and β-actin (bottom panel) antibodies. On the right, molecular weights and the positions of endogenous (*) and GFP-fused STAT1 (**) are indicated.

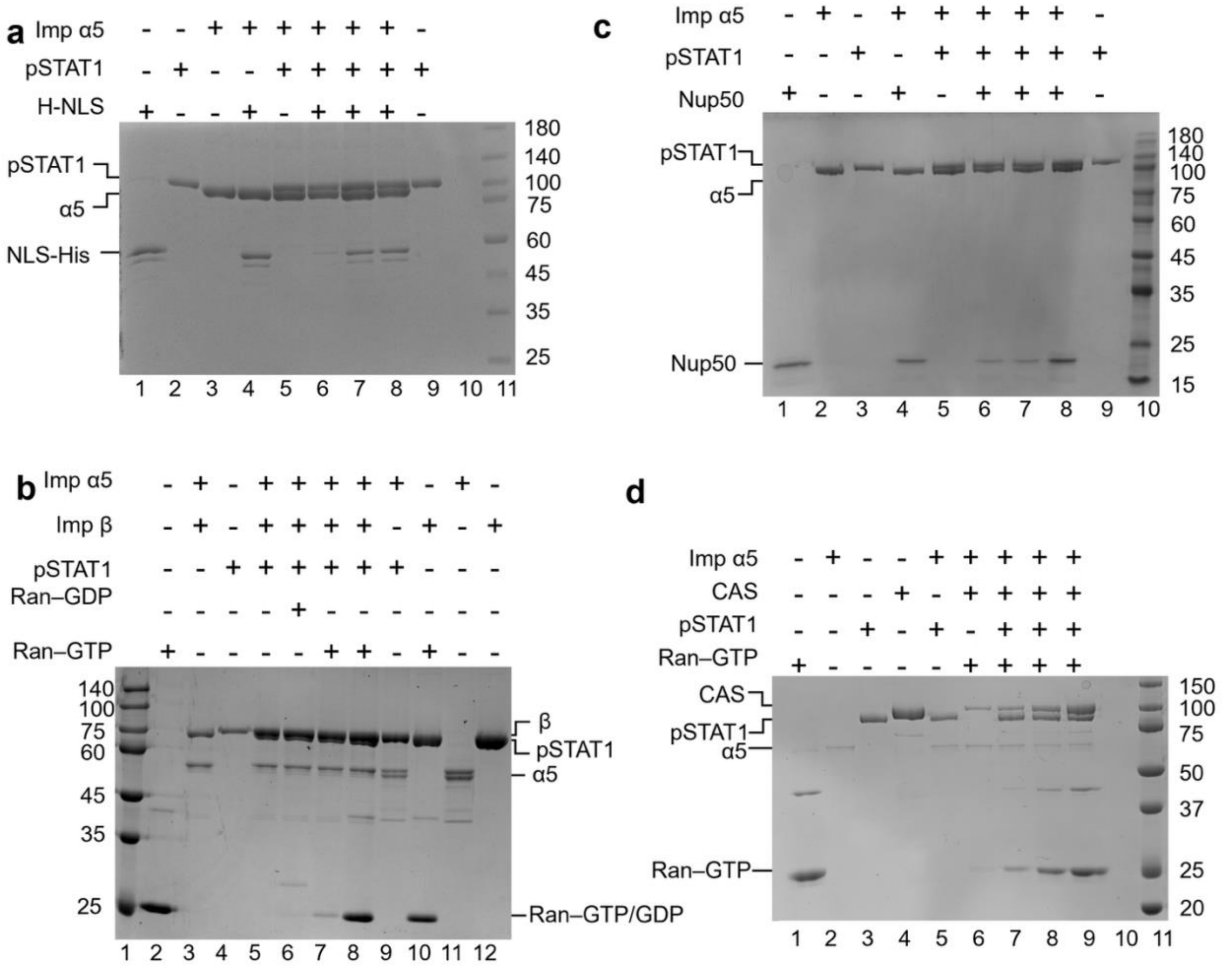

**Supplementary Fig. 5. SDS–PAGE analysis of purified proteins used in native PAGE experiments.** Panels show proteins used in **Fig. 5e** (**a**), **Fig. 5g** (**b**), **Fig. 6c** (**c**), and **Fig. 9a** (**d**). Each gel contains the same protein amounts as loaded in the corresponding native gels shown in the main figures.

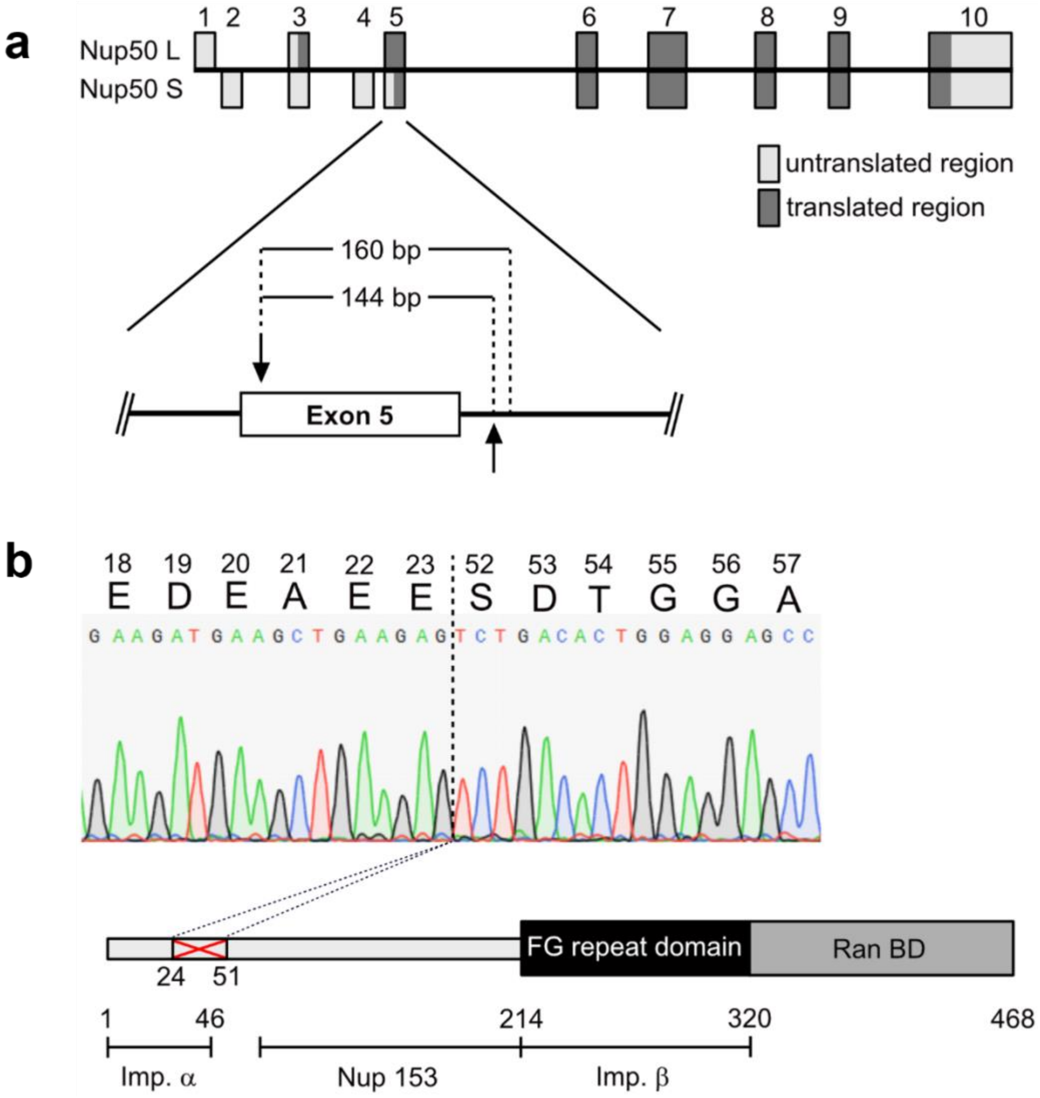

**Supplementary Fig. 6. A 293T cell line expressing Δ24–51 mutant Nup50.** (**a**) Diagram of the genomic structure of human Nup50 transcript variant 2 (Nup50L; NM_007172.4) and variant 3 (Nup50S; NM_153645.2; encoding a rare splice variant that lacks residues 1–28 of Nup50L). Numbers above boxes indicate the exons. The magnified view shows exon 5 and surrounding regions; arrows indicate the regions targeted by the sgRNAs. Sanger sequencing of genomic DNA from the edited cell line revealed deletions of 144 bp and 160 bp. (**b**) Diagram of the Nup50L isoform. Importin α5 binding segment (1–46 aa), Nup153 and importin binding regions, FG repeats, and Ran binding domain (RBD) are indicated. Red X denotes the internal deletion (Δ24–51) in the importin α5 binding segment identified by Sanger sequencing of cDNA (shown above the diagram) from the gene-edited 293T clonal cell line used in this paper. The dotted line on the sequencing chromatogram indicates the site of deletion.

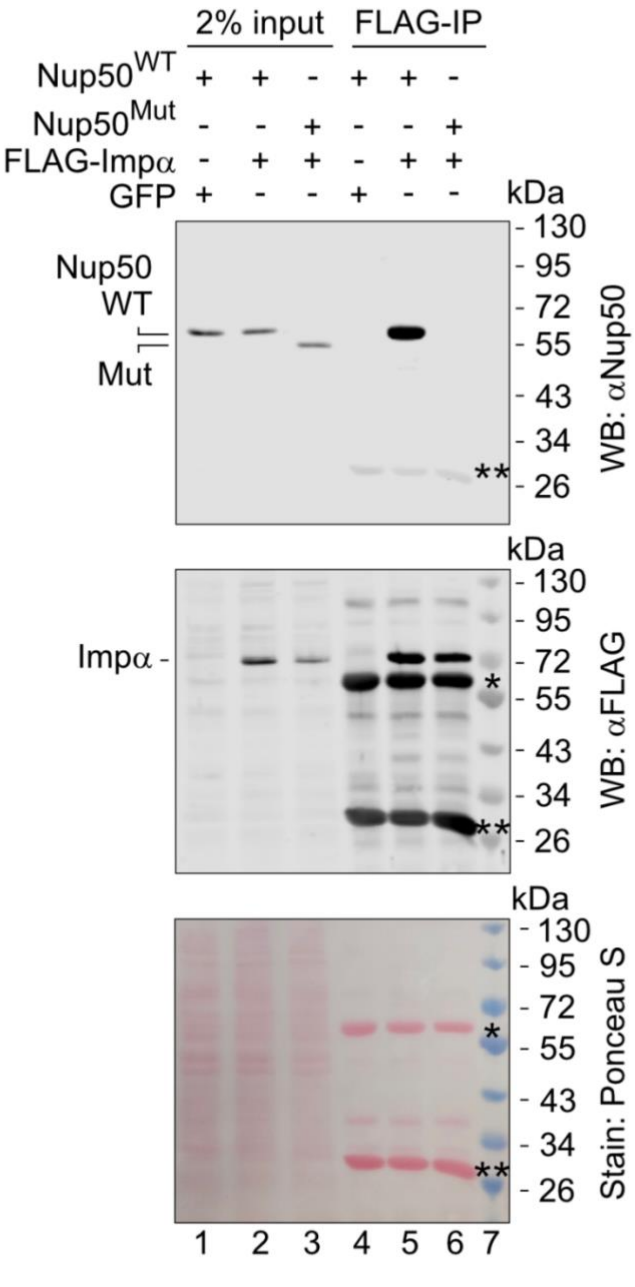

**Supplementary Fig. 7. Binding of WT and Δ24–51 mutant Nup50 to importin α5.** 293T wild-type cells ($Nup50^{WT}$) or gene-edited cells expressing Δ24–51 Nup50 ($Nup50^{Mut}$) were transfected with FLAG-tagged importin α5 or GFP as indicated. Immunoprecipitations (lanes 4–6) were performed from whole cell lysates (lanes 1–3) using anti-FLAG antibody. Shown are Western blots concurrently probed with anti-Nup50 (top panel) and anti-FLAG antibodies (middle panel). Ponceau S staining prior to Western blotting shows total protein loading (bottom panel). Immunoglobulin heavy and light chain positions are indicated by * and **, respectively. Pre-stained molecular weight standards are shown in lane 7.

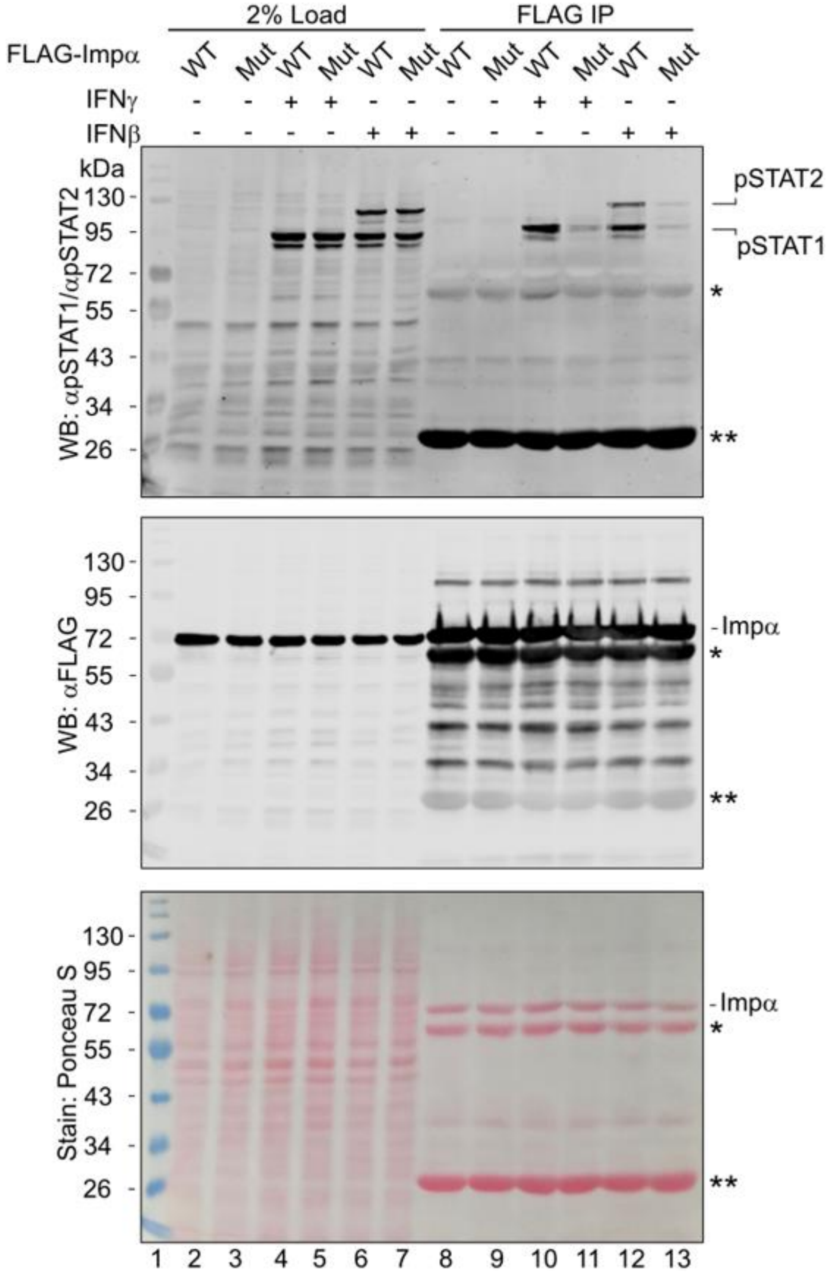

**Supplementary Fig. 8. Binding of activated STAT1 and STAT2 to S1B-mutant importin α5.** 293T cells were transfected with FLAG-tagged importin α5 WT or the Tyr476Ala mutant as indicated. Cells were left untreated or treated for 1 h with IFN-γ or IFN-ß, before immunoprecipitations (lanes 8–13) were performed from whole-cell lysates (lanes 2–7) using an anti-FLAG antibody. Shown are Western blots, probed concurrently with anti-Tyr701-phosphorylated STAT1 (top panel) and anti-FLAG (middle panel) antibodies first, followed by anti-Tyr690-phosphorylated STAT2 (top panel). Ponceau S staining prior to Western blotting shows total protein loading (bottom panel). Immunoglobulin heavy and light chain positions are indicated by * and **, respectively. Pre-stained molecular weight standards are shown in lane 1.

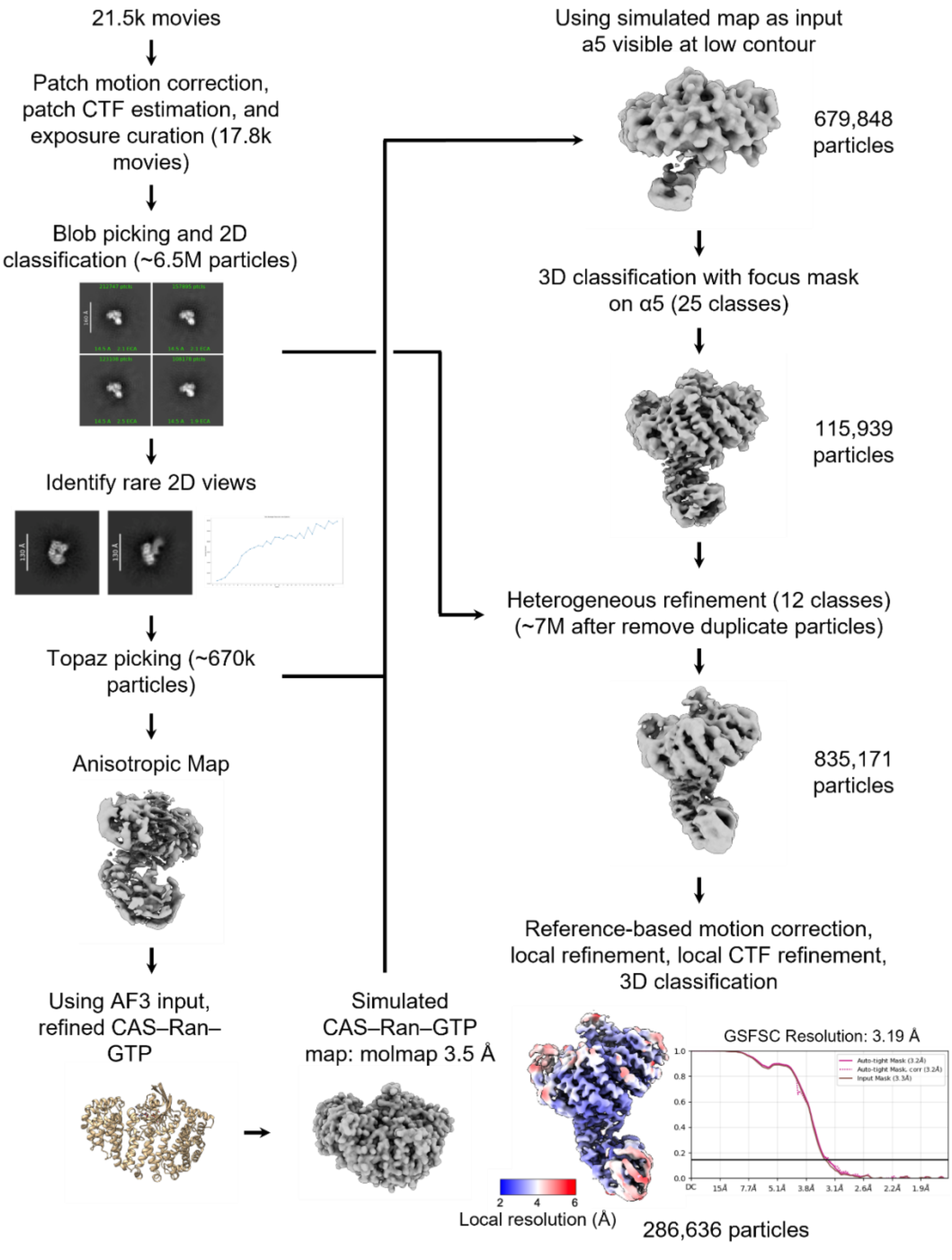

**Supplementary Fig. 9. Workflow of cryo-EM single-particle analysis for the CAS–Ran–GTP–α5 complex.** The bottom-right panels display the local resolution of the final map, color-coded from 2 Å (blue) to 6 Å (red), and the final Fourier Shell Correlation curve. We estimated the resolution using the FSC = 0.143 criterion with independent masking procedures.

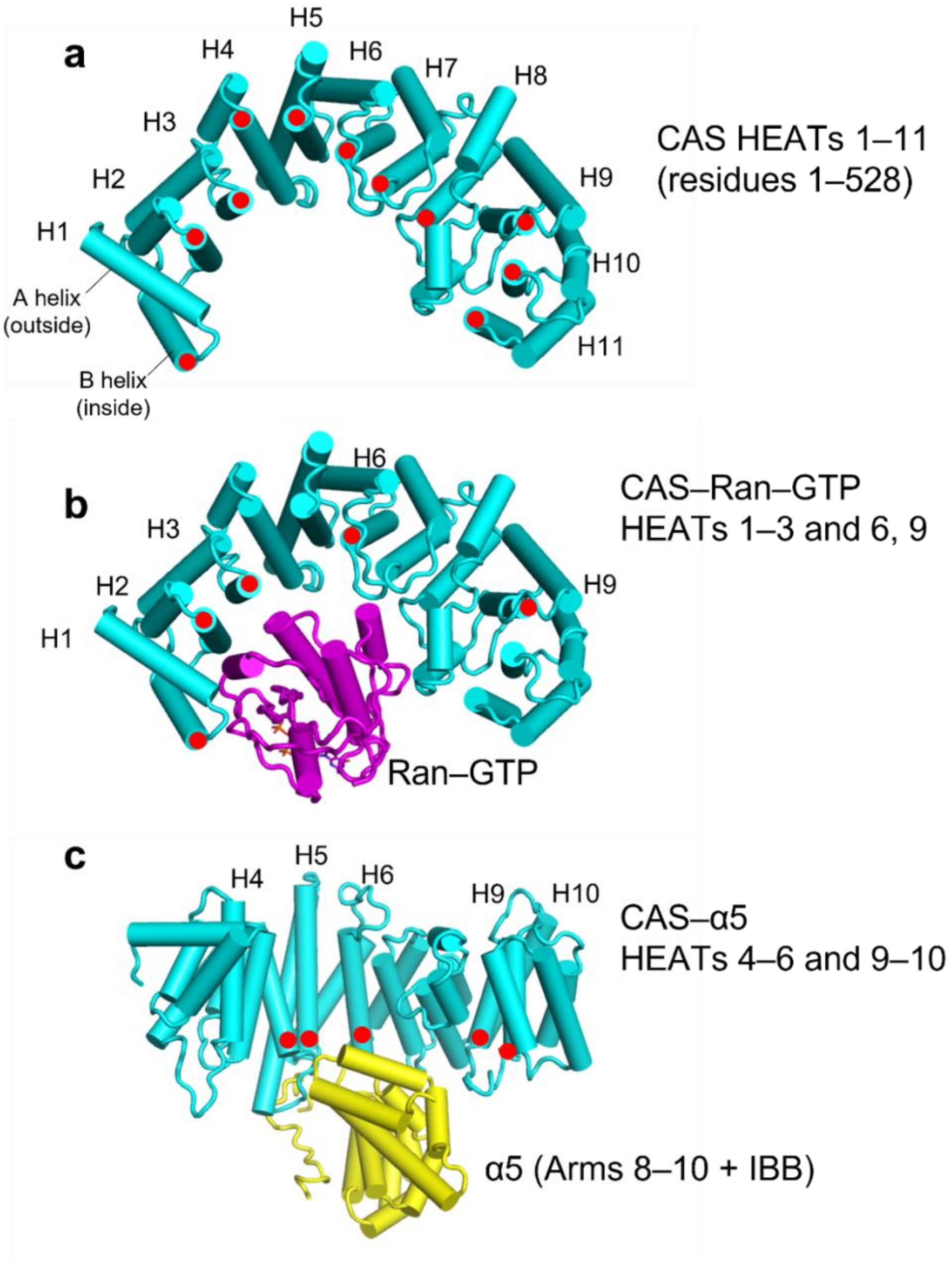

**Supplementary Fig. 10. CAS tertiary structure and its association with Ran–GTP and importin α5.** (**a**) Ribbon representation of CAS residues 1–528 reconstructed in complex with Ran–GTP and importin α5 (only CAS is shown). The reconstruction resolves HEAT repeats H1–H10, with weaker density for H11, corresponding to residues 1–528, each marked by a red dot. (**b**) CAS interacts with Ran–GTP through HEAT repeats 1–3, 6, and 9 (indicated by red dots). CAS is shown in cyan and Ran–GTP in magenta. (**c**) CAS engages importin α5 (yellow) through HEAT repeats 4–6 and 9–10 (indicated by red dots). For clarity, CAS is rotated by 90° relative to panels (**a**) and (**b**).

**Supplementary Table 1. Summary of bonding interactions at protein–protein interfaces**

| pSTAT1–α5 complex (PDB 38YK) | | | | |
|---|---|---|---|---|
| **Interface** | **Salt Bridges (n)** | **Hydrogen Bonds (n)** | **van der Waals Contacts (n) *** | **$\Delta G_{int}$ ** (kcal·mol⁻¹)** |
| **$DBD_T$:α5** | **2**<br>K410:E473<br>K413:E507 | **-** | **5**<br>G384:E474<br>Q412:I465<br>K413:I465<br>A415:G464<br>A415:T463 | –5.3 |
| **$ND_T$:α5** | **2**<br>D49:K480<br>D49:K426 | **2**<br>D42:Y476<br>S51:K480 | **4**<br>H45:Y476<br>S51:T433<br>F52:Y476<br>I55:K480 | –5.5 |
| **$DBD_L$:α5** | – | **2**<br>N381:H487<br>K413:S486 | **14**<br>N381:Q485<br>N381:Y493<br>I382:Y493<br>L383:Y493<br>G384:Q494<br>G384:F497<br>H386:Q494<br>H386:F497<br>K410:Q485<br>E411:S486<br>Q412:E488<br>Q412:S486<br>K413:F483<br>K413:H487 | –2.3 |
| Total bonds | **4** | **4** | **23** | |
| **CAS–Ran–GTP–α5 complex (PDB 11PH)** | | | | |
| **Interface** | **Salt Bridges (n)** | **Hydrogen Bonds (n)** | **van der Waals Contacts (n)** | **$\Delta G_{int}$ * (kcal·mol⁻¹)** |
| **CAS–α5 (S1B)** | **2**<br>K281:D436<br>R384:D479 | **5**<br>H162:E488<br>H162:N489<br>R391:Y476<br>S439:Y476<br>Q444:E473 | **10**<br>K158:E488<br>F164:I492<br>F164:Q441<br>Y220:M435<br>F224:M435<br>L277:M435 | –3.3 |

| | | | | |
|---|---|---|---|---|
| | | | Q280:V434<br>E284:E393<br>Q442:E393<br>T443:E474 | |
| **CAS–α5 (IBB)** | **3**<br>D226:R36<br>E229:K37<br>D233:K37 | – | **6**<br>N167:L33<br>W170:Q34<br>T171:L33<br>K174:L35<br>L227:L35<br>P228:L35 | –1.2 |
| **Ran–GTP–α5** | **1**<br>R95:E501 | **2**<br>R95:F497<br>K132:T506 | **4**<br>R95:I500<br>K99:Y493<br>K99:F497<br>S135:E482 | –1 |
| Total bonds | **6** | **7** | **20** | |

* Van der Waals contacts were identified from atomic coordinates using the PDBsum web server, with an interatomic distance cutoff of ≤3.9–4.0 Å.

** $\Delta G_{int}$ is the free energy of formation of a macromolecular interface upon complex assembly determined by PISA.
